# Paleogenomic insight into the collapse, recovery, and management of American bison

**DOI:** 10.64898/2025.12.24.696034

**Authors:** Jonas Oppenheimer, Joshua D. Kapp, Molly Cassatt-Johnstone, Samuel Sacco, William E. Seligmann, Holland C. Conwell, Sarah Ford, Cassandra Gunn, Lael D. Barlow, Amy Phillips, Kenneth P. Cannon, Lawrence C. Todd, Spencer R. Pelton, Glen MacKay, Kyle Forsythe, Mark A. Edwards, Mark C. Ball, David R.W. Bruinsma, Jessica Z. Metcalfe, John W. Ives, Robert J. Losey, Tatiana Nomokonova, Tomasin Playford, Chris Widga, Craig M. Lee, Karsten Heuer, Wes Olson, Paul Stothard, John Southon, Donalee M. Deck, Christopher N. Jass, Richard E. Green, Lee C. Jones, Gregg P. Adams, Todd K. Shury, Gregory A. Wilson, Beth Shapiro

## Abstract

American bison were pushed to the brink of extinction by the 20th century. This bottleneck and the fragmented nature of remnant populations pose challenges to their resilience, as does human-facilitated admixture between bison subspecies and with cattle. To contextualize current diversity, we sequenced 115 ancient and 45 modern bison genomes from across North America dating back within the last ∼20,000 years. Past bison populations were highly connected, in contrast to structured modern herds. Modern wood bison carry plains bison ancestry from 1920s translocations, while many bison lack cattle ancestry that has previously been believed to be ubiquitous. Our findings reveal the legacy of human impacts on bison in the context of modern conservation and highlight the applicability of ancient DNA for guiding wildlife restoration.

## Main Text

Anthropogenic pressures, including hunting, disease introduction, habitat destruction, and U.S. governmental eradication efforts, nearly drove American bison (*Bison bison*) to extinction by the beginning of the 20th century, with tens of millions of bison being reduced to a few hundred in decades (*1–3*). Concerted efforts by individuals, communities, and governmental entities brought this species back from the brink, with all living bison descended from small populations that survived in managed herds and in the wild (*4–6*). These efforts represented one of the first modern conservation campaigns and fostered the demographic recovery of North American bison, which now number in the hundreds of thousands (*7*).

Management of bison during this 20th century recovery process increased their isolation across small, disconnected herds and also facilitated gene flow between bison subspecies and between bison and cattle (*8*). North American bison include both wood bison (*B. b. athabascae*) and plains bison (*B. b. bison*), which are distinguishable ecologically, phenotypically, and genetically, although morphological boundaries between subspecies are fluid (*9–12*). Wood bison ranged historically across the boreal forest of northern Canada and Alaska (*13*, *14*) and now exist in several small herds (*15*). In the 1920s, the Canadian government translocated ∼7,000 plains bison into Wood Buffalo National Park, at the time home to the last wood bison population of around 1,500 animals (*15*). This translocation makes it possible that wood bison possess some ancestry derived from plains bison. Wood bison are listed as Threatened under Canada’s Species at Risk Act, in part due to introgression with plains bison and the potential loss of genetic diversity from restricted population sizes (*15*).

Cattle introgression is another ostensible genomic legacy of managing post-bottleneck bison populations. During the 20th century, private herd managers conducted breeding experiments aimed at creating bison-cattle hybrids for agriculture (*16*, *17*). While these efforts were generally unsuccessful, it is likely that some bison today possess cattle ancestry due to these efforts (*18*, *19*), and it has recently been suggested that all modern bison carry cattle alleles (*20*). Such domestic cattle ancestry potentially has negative fitness effects (*21*).

Although the history of North American bison stands as a major conservation success, challenges remain (*22*). The genomic effects of the drastic 19th century decline, subsequent recovery, and intervening management of bison populations are poorly understood (*8*). The overwhelming majority of bison today exist in privately-owned production herds, with only a fraction of bison managed in herds specifically for conservation (*23*). Conservation herds tend to be small and geographically isolated, resulting in a fragmented metapopulation occupying a fraction of the range that bison historically inhabited (*24*). Fragmentation risks the loss of overall genetic diversity through the increased effect of drift within subpopulations, as well as stochastic subpopulation extirpation due to disease or climatic events (*25*). Furthermore, the extent of recent admixture between the two subspecies is unknown, and the discovery of wood bison populations free of plains bison introgression would have major ramifications for the conservation of the subspecies. The overall genomic proportion and distribution of cattle ancestry across herds also remains unclear. Understanding cattle introgression dynamics is necessary for guiding movement between bison herds managed as a metapopulation to maintain genetic diversity while minimizing the spread of cattle ancestry (*26*, *27*).

The impacts of the complex recent demographic history bison experienced are difficult to characterize using modern genomes alone (*28*). Recent drift resulting from post-bottleneck isolation of bison herds may obscure prior patterns of population structure and local adaptation and complicate understanding changes in genetic diversity over this timeframe. Determining the nature of recent gene flow between wood and plains bison or between bison and cattle requires unadmixed individuals to serve as a baseline. These baselines can be provided by bison that existed, for example, before the plains bison translocation into Wood Buffalo, or before cattle were introduced to North America. Contemporary conservation has also been hampered by existing molecular tools, such as microsatellites (*29*) and mitochondrial genomes (*30*), which offer limited insight into population history (*31*, *32*). No survey of extant bison diversity across conservation herds has been conducted from a genome-wide perspective. Whole genome ancient DNA presents new opportunities to understand the genomic legacy of these demographic and management events.

In this work, we generated a dataset of 160 bison genomes from across North America spanning the last ∼20,000 years, which were analyzed alongside published genomes from 52 modern bison. We use these data to describe the evolutionary history of North American bison since the Last Glacial Maximum and document changes in population structure and diversity. These genomes provided a reference for characterizing recent, human-facilitated admixture between bison subspecies and between bison and cattle. This information can be used to guide decision making to support the continued restoration of the American bison as an ecologically, culturally, and economically vital species.

## Results

We define “modern” as bison that lived within the last ∼100 years, whereas “ancient” refers to bison that lived before and during their recent population collapse. We sequenced 115 ancient and 45 modern bison genomes from across North America dating back to the Last Glacial Maximum to a median of 1.75-fold coverage (0.01- to 34.43-fold; Fig. 1A and table S1). We added 52 publicly available modern bison genomes (including 6 bison dating between 1886-1937) into our dataset [table S2; (*20*, *33–39*)], for a total of 212 individual bison from the past ∼20,000 years. A map of modern herds sampled in this study is presented in fig. S1. We also generated new radiocarbon dates for 73 of the sequenced bison specimens which had not been previously dated (tables S1 and S3).

**Figure 1:**
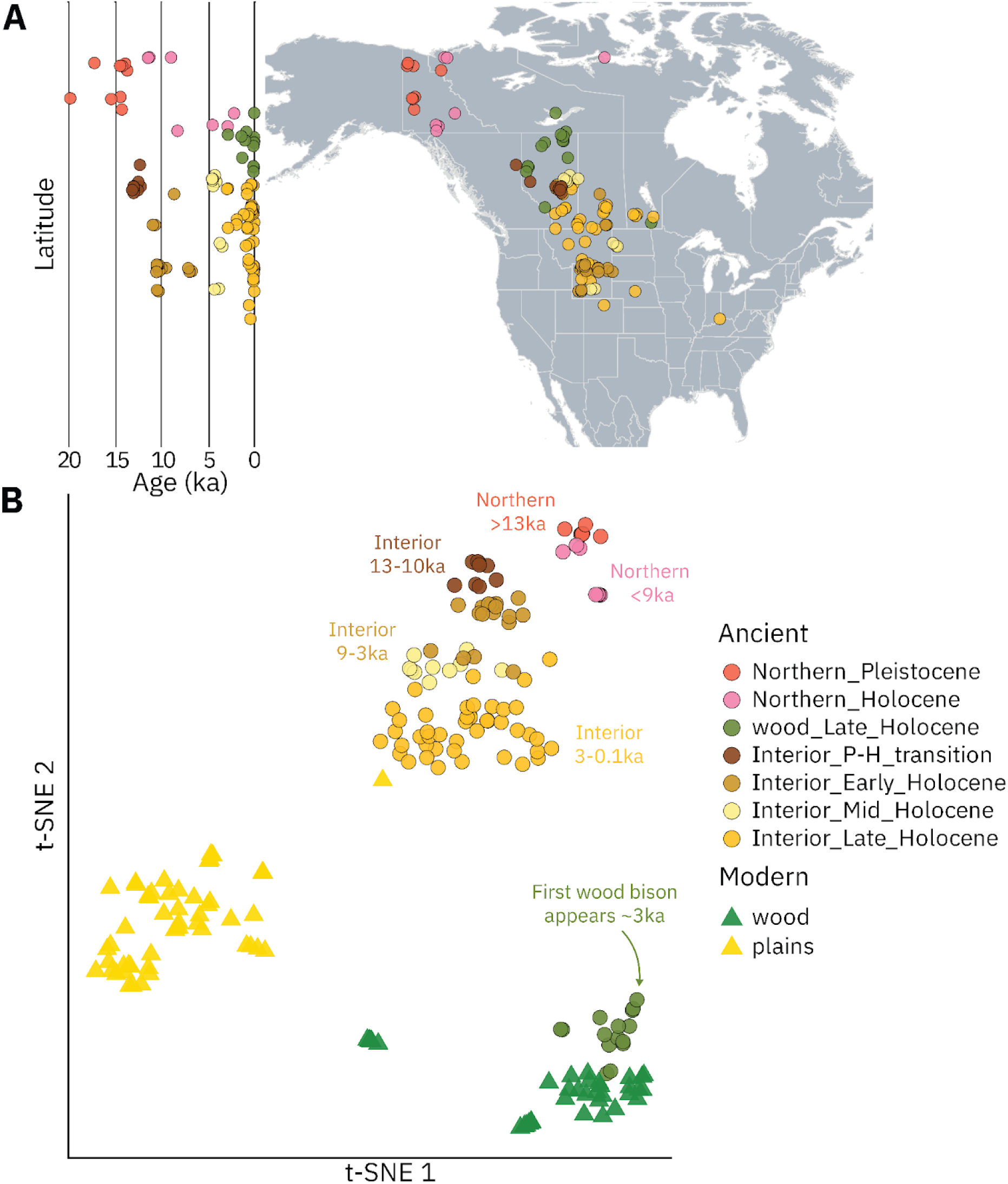
Sampling and genetic structuring of North American bison after the Last Glacial Maximum. (**A**) Map showing location and age of all sampled ancient bison. Jitter has been added to sample coordinates for visualization. (**B**) f4-statistic-based t-SNE analysis showing genetic clustering of ancient and modern North American bison. Samples are colored based on temporal, geographic, and genetic groupings, corresponding to three main groups: remnant northern steppe bison (red and pink), wood bison (green), and Interior/plains bison (brown and gold).

**Figure 2:**
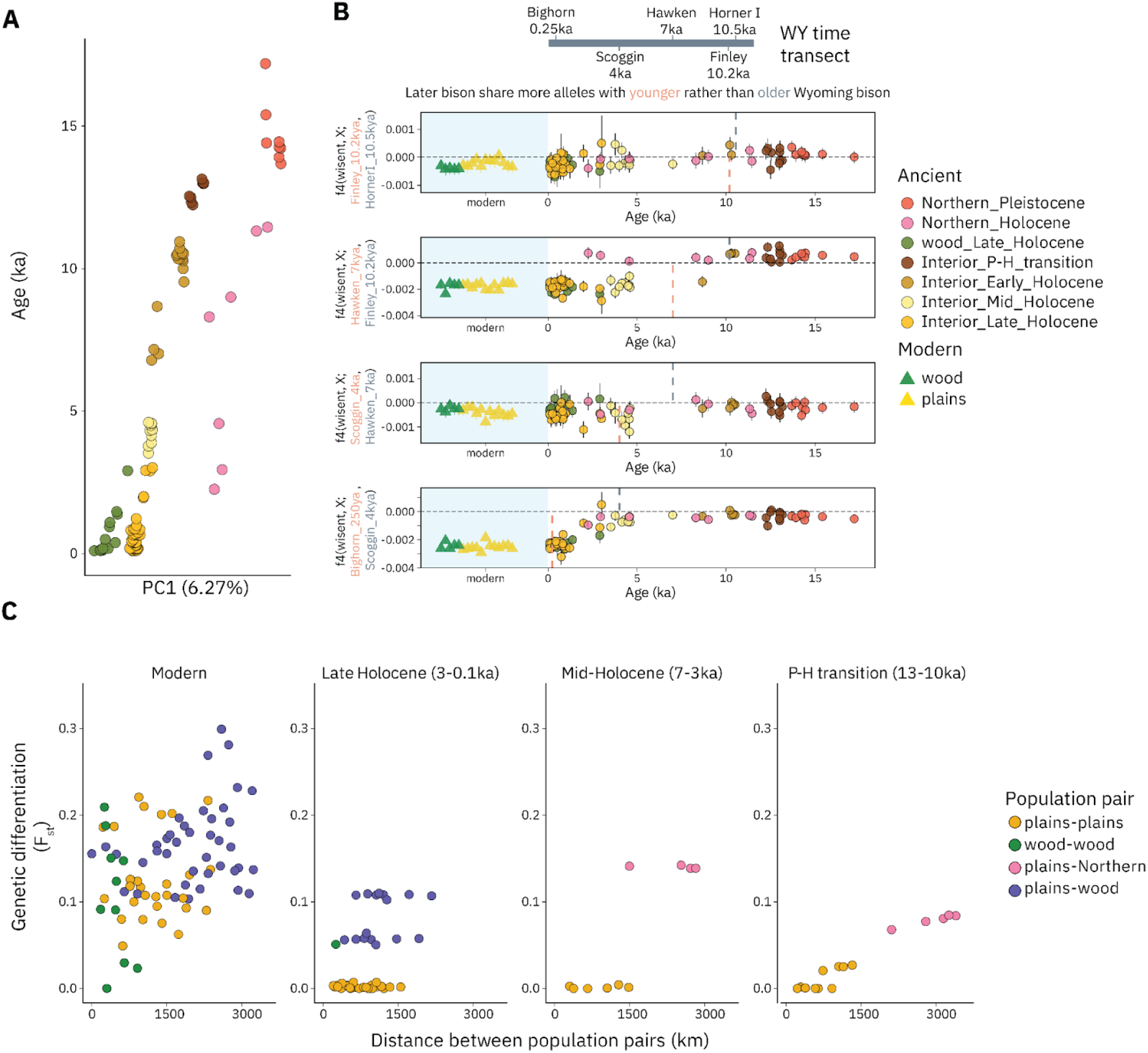
Extensive connectivity in North American bison throughout the Holocene. (***A***) Age by PC1 from an f4-statistic-based PCA computed using all ancient North American bison. (**B**) Relative allele sharing of all bison to pairs of sites using a time transect of bison from Wyoming spanning 10,000 years. Five sites are shown, grouping the Bighorn Basin samples as a single site, and each is compared to the sites with dates that occur immediately before and after. The gray line shows the date of the older site in the comparison, while the pink line shows the date of the younger site. Lower f4-statistic values indicate more allele sharing with the younger site, relative to the older one. Bars show two standard errors. (**C)** F_st_ between pairs of bison populations throughout the Holocene, split into four time periods. Each point represents a comparison between two population pairs, which were grouped based on geographic and temporal similarity. Colors correspond to the populations in the comparison.

As morphological distinctions between subspecies are imprecise, subspecies identification of fossils is challenging. The geographic distribution of genetically determined wood bison samples refines our understanding of their former range (*13*). Areas that were previously thought to represent wood/plains hybrid zones (Fig. 4A), such as north-central Alberta south of Wood Buffalo, yielded only plains bison in our sampling (n=16; Fig. 1A). However, wood bison were also present in unexpected locations, including in western Alberta near Banff and in southern Manitoba at The Forks National Historic Site, both of which are further south and are located further west and east, respectively, than the previously supposed historical wood bison habitat (*43*). It is possible that wood bison were widely distributed throughout the Canadian boreal forest and not confined to its northern reaches, although inferences of wood bison distribution from genetic data are potentially complicated by trade (*44*) or long-range human (*45*, *46*) or animal (*47*, *48*) mobility.

**Figure 3.**
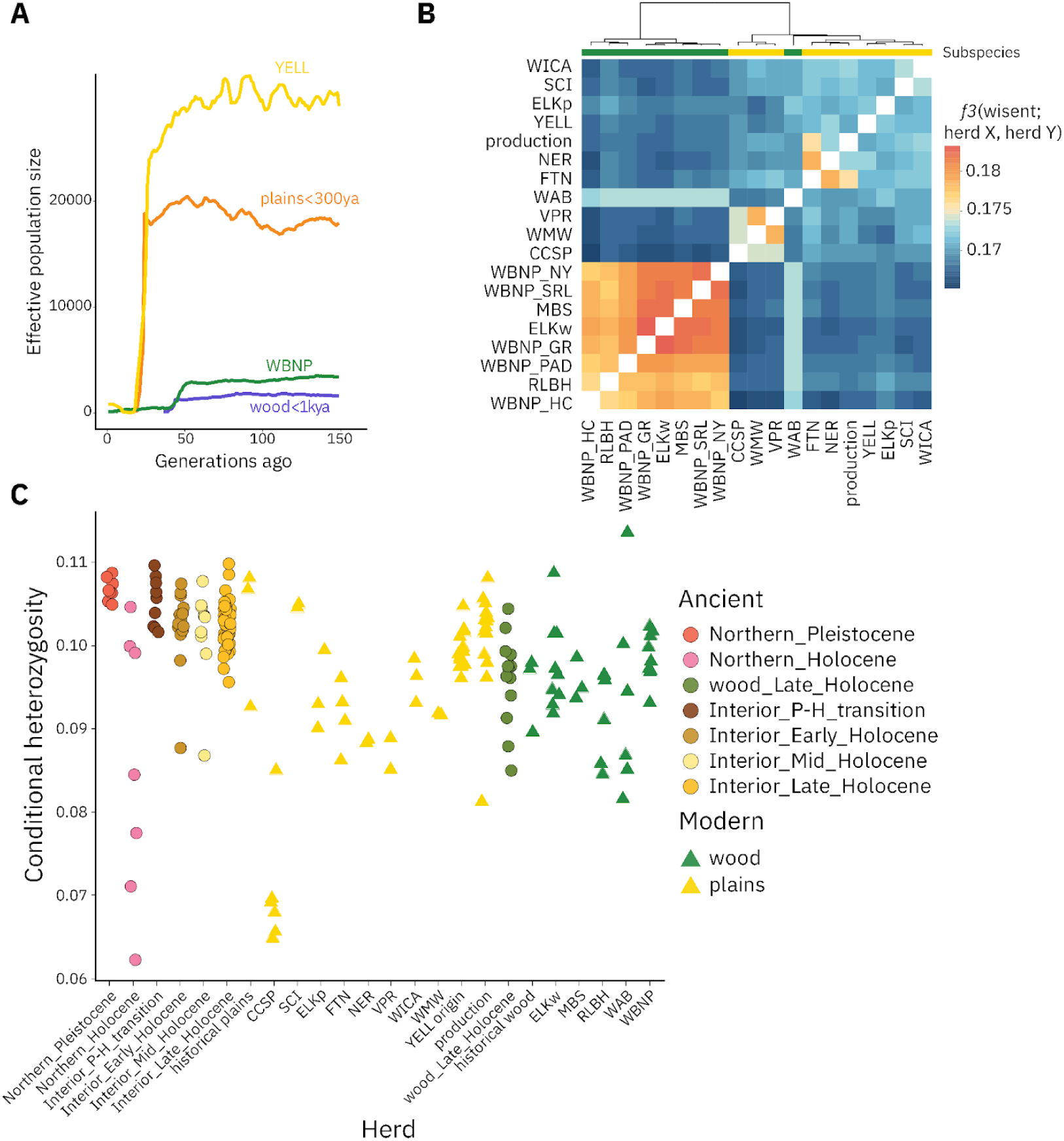
Rapid increase in genetic differentiation without loss of diversity in modern bison. **(A)** Effective population size reconstructions for modern Yellowstone (n=15) and Wood Buffalo (n=9) National Park herds, along with ancient plains (<300ya; n=18) and wood (<1ka; n=11) bison. Ancient curves are shifted back to match modern curves. (**B)** Heatmap of outgroup f3-statistics and allele sharing-based hierarchical clustering among modern bison herds. (**C)** Conditional heterozygosity, measured as the mismatch rate of two randomly sampled reads across a set of pre-ascertained variants, for modern herds and ancient populations. Herd abbreviations: CCSP = Caprock Canyons State Park; SCI = Santa Catalina Island; ELK = Elk Island National Park; NER = National Elk Refuge/Grand Teton National Park; FTN = Fort Niobrara National Wildlife Refuge; VPR = Vermejo Park Ranch; WICA = Wind Cave National Park; WMW = Wichita Mountain Wildlife Refuge; YELL = Yellowstone National Park; MBS = Mackenzie Bison Sanctuary; RLBH = Ronald Lake Bison Herd; WAB = Wabasca; WBNP = Wood Buffalo National Park. The WBNP herd is split into geographical subpopulations: GR = Garden River; SRL = Slave River Lowlands; PAD = Delta; NY = Nyarling; HC = Hay Camp.

**Figure 4.**
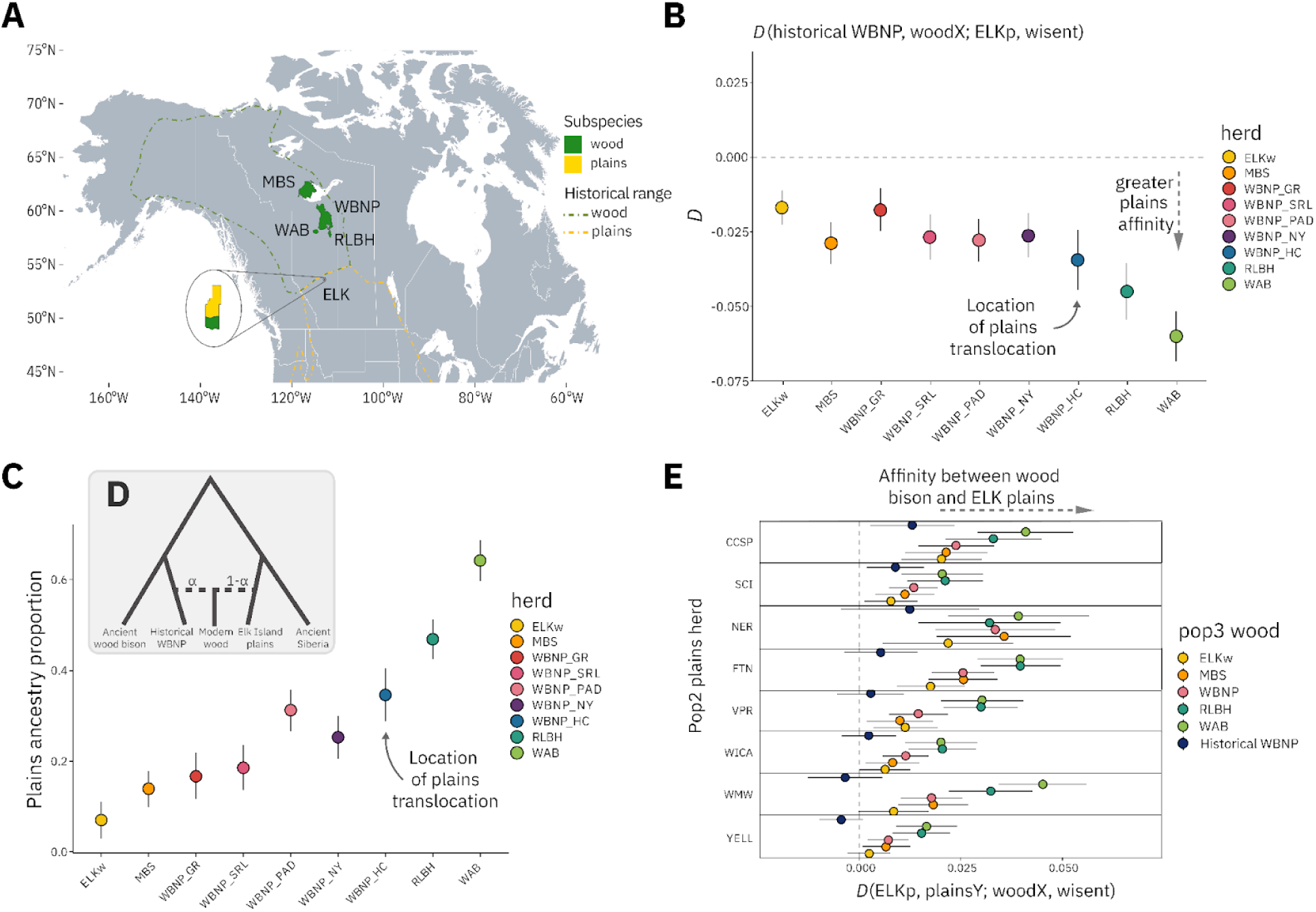
Widespread though variable plains bison ancestry in all modern wood bison. (**A**) Locations of modern wood bison herds and their previously estimated historic range. Elk Island National Park contains physically separated plains and wood herds. **(B)** Allele sharing between modern wood bison herds and Elk Island plains bison, relative to historical Wood Buffalo National Park samples. The Hay Camp subpopulation is highlighted as being closest to the geographic origin of the plains bison translocations into Wood Buffalo. **(C**) Estimate of plains bison ancestry proportions across modern wood bison populations and subpopulations, performed using f4-ratio outlined in (**D)**. (**E)** Relative allele sharing between modern and historical wood bison to either the Elk Island plains herd or other modern plains herds. Each row represents a different modern plains herd used in the comparison. Error bars represent ±3 standard errors. Herd abbreviations are the same as those in Fig. 3.

Bison were stratified by time along the second t-SNE axis, as Late Pleistocene and Early Holocene individuals occurred in the upper portion, while those with more recent dates appeared lower. Contemporaneous bison grouped together even when at opposite geographic extremes of our sampling (Fig. 1A), which spans the majority of bison distribution since the last Ice Age, encompassing the Interior Plains and its margins along the Canadian Shield and Intermountain West and Canadian Intermountain Region (*49*, *50*). Some geographic substructuring was present in remnant northern bison from the Yukon and Northwest Territories, which were shifted upward on the second axis (Northern_Pleistocene and Northern_Holocene; Fig. 1B), possibly because they possess additional Siberian-associated ancestry not present in post-Last Glacial Maximum Interior bison [fig. S4; (*51*)]. Holocene northern bison were shifted towards Interior bison relative to those from the Late Pleistocene, likely as a result of a northward expansion of Interior bison after the opening of the Ice Free Corridor (*52*). All pre-collapse Holocene Interior bison formed a distinct cluster, also stratified temporally (fig. S6). These results demonstrate strong population connectivity in bison across North America over the past 20,000 years, with minimal genetic structuring over vast geographic distances, particularly since the opening of the Ice Free Corridor ∼13ka (*53*, *54*). Modern bison, including both wood and plains subspecies, grouped separately from pre-bottleneck bison populations, likely due to increased drift associated with the population collapse and subsequent isolation during and after demographic recovery (*26*).

## Extensive connectivity over the Holocene

Examining bison phylogeography over the last several thousand years enables direct reconstruction of how bison existed across the landscape while contextualizing recent anthropogenic impacts on modern herds. Our *f-*statistics-based dimensionality reduction analyses revealed that time was a primary factor separating bison populations after the Last Glacial Maximum (Fig. 1B). In a principal component analysis (PCA) of all ancient North American bison, PC1 was strongly associated with sample age (Pearson’s *r*_age,PC1_ = 0.86, *P* = 7.84 x 10^-34^; Fig. 2A). Temporal structuring was most obvious within Interior bison, with distinct clusters forming for wood bison and remnant northern bison, although within these groups similar temporal gradients were observed (fig. S6). This pattern suggests widespread gene flow through time between most Holocene North American bison, regardless of geography.

To understand this apparent connectivity of Interior bison in greater detail, we assembled a 10,000 year time transect, which comprised 16 bison from five archaeological sites in Wyoming (*55*) along with 11 pre-bottleneck bison (>180ya) from throughout Wyoming’s Bighorn Basin. We then examined how connections between Wyoming and the rest of North America shifted through time. Specifically, we tested for differential allele sharing between pairs of temporally adjacent Wyoming sites to all other bison in our dataset. When we contrasted relative allele sharing for non-Wyoming bison to a series of progressively younger pairs of Wyoming sites adjacent in age, bison which postdated both sites generally had more affinity with bison from the younger site than the older site, particularly for Interior bison (Fig. 2B). This is consistent with extensive gene flow connecting bison from these Wyoming sites and all other bison throughout the Holocene. Such connectivity suggests high dispersal over this period, although isotopic evidence attests to limited extreme long-range movement (*47*).

In contrast to the widespread connectivity observed throughout much of the Holocene, present-day bison exhibit much stronger population structure. Modern bison clustered by herd origin in t-SNE space (Fig. 1B), indicating that the management of bison in small, isolated conservation herds has increased genetic differences among herds, likely through drift and possibly heightened by management practices (*56*). We compared the relationship between geographic distance and genetic differentiation, as measured by F_st,_ throughout different periods since the end of the Pleistocene, grouping ancient bison by time and geography and modern bison by herd (Fig. 2C). We found almost no genetic differentiation in all pre-bottleneck comparisons between pairs of Interior bison populations (median F_st_ = 0.001; range: 0-0.011), even when separated by thousands of kilometers. Northern and Interior groups did have larger differentiation (median F_st_ = 0.109; range: 0.062-0.142) which grew over time, perhaps driven by a loss of diversity in northern populations throughout the Holocene (Fig. 3C). Ancient wood bison had higher levels of differentiation from Interior bison populations, relative to within-Interior comparisons, after their emergence in the Mid/Late Holocene.

Though geographic distances between modern populations are mostly the product of human management, confounding any relationship between genetic and geographic distance, genetic differentiation between any pair of modern plains herds was generally greater than that found between Interior and northern bison in the Early and Mid-Holocene, reflecting isolation and loss of connectivity in modern herds. Differentiation between modern wood and plains bison was also increased in present-day bison, exceeding that between any pair of North American bison populations after the Last Glacial Maximum and comparable to that between North American and Eurasian steppe bison in the Pleistocene (F_st_ = 0.218±0.001).

## Effects of a drastic population decline

Demographic reconstructions of both plains and wood bison using GONE2 (*57*) reveal the timing and magnitude of their recent population collapse (Fig. 3A). In agreement with historical sources (*1*), we find that bison experienced a profound decline in effective population size (*Ne*) within the last few centuries. For the Yellowstone herd (n=15), this bottleneck began around 1775-1850 (25 generations ago), assuming a generation time of 7-10 years (*58*), and reached a nadir by 1875-1920 (15 generations ago), with a reduction in effective population size of about three orders of magnitude (*Ne* of 46 vs. 31,214). Using a larger-scale geographic sample of ancient Interior bison dating within the last 300 years (n=18) — the most recent ancestors of modern plains bison — we recovered a similar decline. Wood Buffalo National Park wood bison (n=9) were inferred to have had much lower pre-bottleneck effective population sizes than plains bison (mean *Ne* of 3,152) and to have also undergone a recent population reduction, though their decline was proportionally less severe (minimum *Ne* of 127) and began nearly 200 years earlier (52 generations ago), possibly coinciding with the expansion of the Canadian fur trade (*59*). As with plains bison, ancient wood bison (n=11) also demonstrate a decrease in *Ne* consistent with that estimated using modern Wood Buffalo bison.

Following the population collapse, several distinct wild or captive foundational herds survived, from which all bison today originate (*20*). Clustering of modern bison herds using pairwise outgroup *f3*-statistics demonstrated a large degree of genetic structure across modern bison (Fig. 3B). Bison were separated first by subspecies, and then formed distinct clusters within each subspecies which were consistent with herd foundation histories (*8*, *20*). Four major groups of plains bison were evident: National Elk Refuge/Grand Teton National Park and Ft. Niobrara National Wildlife Refuge; Caprock, Vermejo, and Wichita Mountains; Yellowstone and Elk Island; and Wind Cave, Catalina, and production herds. This clustering by outgroup *f3*-statistics shows that substantial drift following the population collapse and subsequent establishment of conservation herds, as well as the continued isolation of these herds, has led to the elevated levels of genetic differentiation observed in modern bison.

Despite its dramatic scale, the 19th century bison population bottleneck appears not to have resulted in a reciprocal loss of genetic diversity (Fig. 3C). Wood bison had slightly lower genetic diversity (median conditional heterozygosity of 0.096) during the Holocene than Interior bison (median of 0.102), consistent with their lower estimated effective population sizes. Estimates of heterozygosity were similar pre- and post-bottleneck for both wood (modern median heterozygosity = 0.097) and plains bison (modern median heterozygosity = 0.099). However, variation in heterozygosity across individuals was somewhat larger in modern individuals (standard deviations of 0.0064 and 0.0114, respectively) relative to ancestral wood and plains bison (standard deviations of 0.0053 and 0.0036), such that these demographic events may have had differing effects on diversity across herds. For modern bison, the proportion of the genome contained in runs of homozygosity (ROH) greater than 1Mb is correlated with estimated heterozygosity (Pearson’s *r*_heterozygosity,% ROH_ = -0.78, *P* = 1.01 x 10^-14^; fig. S7), suggesting that the greater variation in diversity observed across modern herds is driven by recent inbreeding. The Caprock Canyons Texas State bison herd, which had the lowest estimated heterozygosity (Fig. 3C) and highest ROH proportion (fig. S7), has displayed phenotypic signs of inbreeding (*60*).

## Historical translocations facilitated subspecies hybridization

All modern wood bison (Fig. 4A) have evidence of plains bison introgression stemming from 1920s translocations of plains bison into Wood Buffalo (*43*). As a baseline for detecting recent hybridization, we used two ∼175ya bison recovered from Wood Buffalo that existed before the historical translocations but had a close genetic relationship with modern Wood Buffalo wood bison (Fig. 1B). *D*-statistics confirmed excess derived allele sharing between plains bison and modern wood bison relative to these pre-1920s Wood Buffalo bison (−14.21 ≤ *Z* ≤ -4.98; Fig. 4B). Similar allele sharing was not detected in ancient wood bison and all ancient wood bison samples formed a clade relative to plains bison (fig. S11), confirming that subspecies admixture is a product of recent human action, rather than an ongoing historical process. After their separation of at least 3ka, it appears there was little recurrent contact between subspecies. A cline of plains bison ancestry is evident across and within wood bison herds, and the Wood Buffalo subpopulations (*61*) with the highest levels of plains ancestry are those geographically closest to the site of the 1920s plains bison translocations (Fig. 4C). Elk Island National Park had the lowest overall proportion of plains ancestry (7±4%), while the Ronald Lake and Wabasca herds had the highest (47±4% and 64±5%, respectively). We estimated plains bison ancestry proportions using the *f4*-ratio depicted in Fig. 4D.

Elk Island contains two separate bison herds, one of each subspecies (Fig. 4A). The Elk Island plains bison herd, derived from the same population of bison translocated to Wood Buffalo in the 1920s (*62*), serves as the best proxy for the source of plains bison ancestry in modern wood bison. Modern wood bison share significantly more alleles with Elk Island plains bison than other plains herds (0.88 ≤ *Z* ≤ 8.37; Fig. 4E), implicating historical translocations as the source of subspecies introgression. The same pattern is not observed in historical pre-translocation Wood Buffalo bison (−1.66 ≤ *Z* ≤ 2.50; Fig. 4E), confirming that this is the result of recent interbreeding, rather than ancestral contact. We cannot rule out bidirectional gene flow (wood bison admixture into Elk Island plains bison), but replacing Elk Island with other plains bison herds yields similarly negative statistics, confirming the presence of plains bison ancestry in modern wood bison (fig. S12). In modeling with the qpWave/qpAdm framework (*63*) using the four major plains bison groups (from Fig. 3B) as source populations, Elk Island is sufficient as a single source to explain plains ancestry in all wood bison populations, although the Wabasca herd is perhaps better fit with an additional non-Elk Island source, which could explain its higher levels of plains ancestry (tables S5-S7).

Finally, we found that the Ronald Lake and Wabasca wood bison herds share a close genetic relationship with other modern wood bison populations. The origins of these herds had been unclear (*15*) and they had previously been shown to be genetically distinct from other wood bison (*11*). This was thought to potentially be due to these herds both deriving from separate ancestral lineages and lacking plains bison ancestry, making them a unique source of wood bison variation. We found that both herds share significantly more alleles with modern wood bison than they do with any ancient wood bison individuals (−21.68 ≤ *Z* ≤ -3.11; fig. S13). This is consistent with a very recent separation from Wood Buffalo bison for both herds, possibly following initial plains-wood gene flow, with subsequent additional admixture with plains bison leading to their higher proportion of plains ancestry.

## Cattle ancestry is not ubiquitous in modern bison

Introgression with cattle has been proposed as a pervasive feature of modern bison population history (*20*). Our large sample of ancient bison that existed from before cattle were introduced to North America served as a secure baseline for assessing recent domestic cattle hybridization.

Using reads mapped to the bison genome (*34*) to mitigate reference bias causing artifactual attraction to cattle, we tested for allele sharing with cattle in 32 modern bison sampled from nine herds (Fig. 5A). This sampling included some bison previously reported to have cattle ancestry detected using microsatellites [L.C. Jones, unpublished data and (*19*)]. Three of these 32 bison had significant allele sharing with cattle as compared to ancient bison (3.35 ≤ *Z* ≤ 4.38; Fig. 5A, figs. S14 and S15) confirming the presence of cattle ancestry in some modern bison. Although this analysis only included samples from a limited number of bison herds, the distribution of these introgressed individuals among three different herds demonstrated that cattle ancestry is not localized to any particular group. Additionally, that many of the bison examined had no detectable excess affinity to cattle suggested that cattle ancestry may be less common than has been believed.

**Figure 5.**
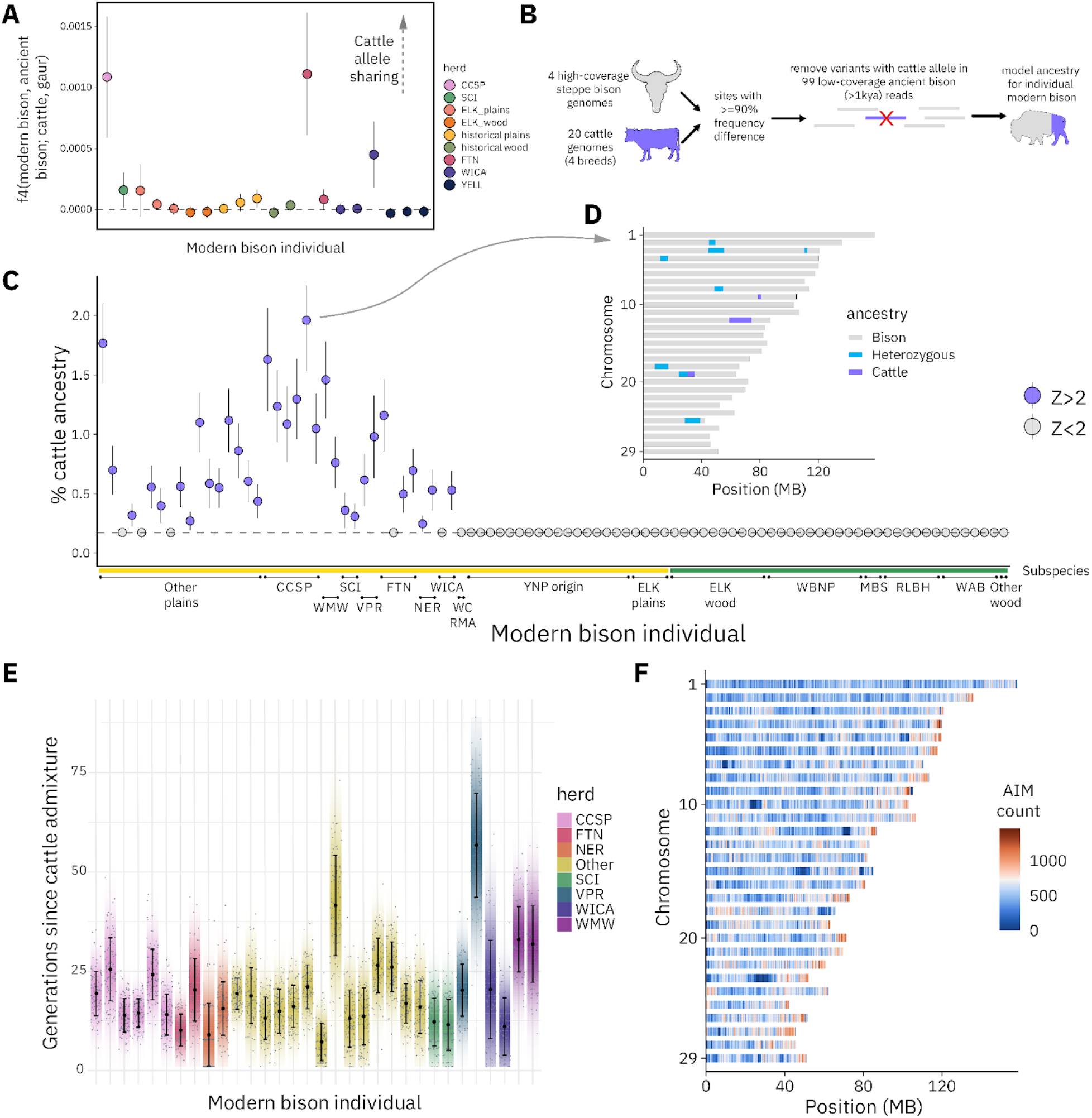
Identifying the amount and location of cattle ancestry in modern bison. (*A*) f4-statistic testing for affinity between modern bison individuals and cattle, relative to pre-cattle introduction bison. Herd origin is shown by the colors. Error bars represent ±2 standard errors. (**B)** Schematic showing the local ancestry inference procedure used to identify segments of cattle ancestry in individual bison genomes. (**C)** Summary of total cattle ancestry proportions for all modern bison individuals. Each point is an estimate for an individual, which are grouped by herd and then subspecies. Yellow bars denote plains bison and green bars denote wood bison. Individuals with detectable proportions of cattle ancestry (Z>2) are shown in purple, those that did not have statistically significant cattle ancestry are shown in gray. Error bars represent ±2 standard errors. **D)** Example of cattle ancestry local ancestry inference of a CCSP individual, which was inferred to have the most total cattle ancestry across all bison examined. **E)** Estimates of cattle admixture timing across sampled bison, colored by herd origin. Gray dots represent bootstrap replicates, while error bars depict one standard deviation and strips portray 95% confidence intervals. **F)** Distribution of bison-cattle ancestry informative marker density in 250kb windows across all autosomes. Herd abbreviations are the same as those in Fig. 3. “Other” refers to bison from non-conservation, largely production, herds.

Considering the surprising lack of evidence for widespread cattle introgression in this initial test, we sought to survey cattle ancestry more broadly across modern bison. We used a local ancestry inference approach (*64*), modeling ancestry within individual bison genomes as coming from one of two sources: an ancient bison panel with four high-coverage (≥17-fold coverage) Beringian steppe bison genomes dating between ∼50-10ka, and a modern taurine cattle panel, which had 20 individuals from four breeds (*36*) that are plausible sources of cattle ancestry in bison (Fig. 5B; table S8). For this analysis, we used a set of ancestry-informative markers (AIMs) with a minimum 90% frequency difference between the two panels; to take advantage of our larger dataset of low-coverage ancient bison genomes, we filtered these sites to remove any where ancient bison (>1ka) had reads carrying the cattle allele, eliminating variation deeply shared between species. We also removed sites variable among cattle [represented by the 1000 Bull Genomes Project Run 9 call set (*65*)] and which were not confidently assigned as heterozygous in “Buzz”, an F1 bison-cattle hybrid (*34*), resulting in a final set of 5.36 million AIMs. For individuals with known ancestry, such as “Buzz”, this method assigned the correct ancestry proportions (prior to subsequently filtering non-confident sites in “Buzz”; fig. S16). Local ancestry inference also provided results consistent with allele sharing statistics, as the three individuals with significant cattle affinity had detectable tracts of cattle ancestry.

Cattle ancestry was present in 32 of the 97 modern bison sequenced (Fig. 5C). Although the majority of bison we examined displayed no evidence of cattle admixture, we note that our sample sizes for several herds are low, consisting only of two individuals, and that our selection of bison was non-random. However, since our sampling was intentionally biased to include several bison suspected of being introgressed through prior microsatellite analysis, we may have overestimated the extent of cattle admixture within and across herds. Nevertheless, most herds we examined had at least one individual free of cattle introgression, and several, including both Elk Island and Yellowstone-origin bison, had no individuals identified as introgressed across somewhat larger sample sizes (n=4 and 16, respectively). Furthermore, no wood bison (n=35) had evidence of cattle ancestry, such that historical cattle admixture may have been limited to plains bison.

We found cattle ancestry to be more prevalent in bison outside of herds managed for species conservation, though we did identify two individuals from production herds without detectable cattle admixture, such that introgression is not ubiquitous in production contexts (Fig. 5C). When present, the levels of introgression were low (0.07 to 1.79%), and tended to occur in large, discrete, heterozygous segments (fig. S17). An example is shown in Fig. 5D from a Caprock Canyons individual, which was identified as possessing the highest proportion of cattle ancestry in our study (1.79±0.21%). The distribution of cattle ancestry is consistent with limited and recent hybridization between bison and cattle. Dating of the hybridization event in admixed individuals suggests that introgression took place ∼20 generations in the past, coinciding with when plains bison reached their nadir (Fig. 5E). There is also some evidence that cattle ancestry is distributed nonrandomly across the genome (Chi-squared *P* = 0.016; fig. S19), although larger sample sizes are likely necessary to determine whether this is due to patterns of recombination, demography, or selection. Together, these results suggest that cattle ancestry is much less common than previously thought and could potentially be lost over time through drift, as long as gene flow between herds is frequent enough to prevent particular segments of cattle ancestry from becoming fixed within herds.

Examining the distribution of ancestry-informative sites between our ancient bison and cattle source panels provides insight into why previous analyses have suggested that all modern bison have cattle ancestry (*20*). The density of sites at high frequency difference between the two panels varied widely across the genome (Fig. 5F). Areas with low differentiation between the two panels (a lack of ancestry-informative sites) were often previously identified as originating from cattle but are not in our analysis [fig. S21; (*20*)]. In some cases, these correspond to areas that are highly repetitive or poorly resolved in the cattle reference genome, leading to poor mapping and a lower density of called sites. The inclusion of ancient bison sampled prior to the introduction of cattle allows for distinguishing between recent introgression and other causes of genomic similarity, including incomplete lineage sorting, genomic conservation, or technical artifacts, and highlights the difficulty in disentangling introgression from other processes. These challenges have likely led to prior overestimation of the prevalence of cattle ancestry both within and across individuals.

## Discussion

The near-extinction of North American bison at the end of the 19th century and their subsequent history of rescue, demographic recovery and management have left an undeniable impact at the genomic level. Modern bison herds, including both wood and plains subspecies, cluster separately from bison that existed prior to this bottleneck. These shifts in genetic structure can be attributed to founder effect and drift associated with their population crash and recovery, during and after which bison were kept in small, isolated groups (*8*), leading to higher genetic differentiation among bison herds today than in the past. Holocene North American bison had strikingly little geographic structuring, with close genetic associations evident between bison at disparate ends of the North American continental interior, indicating high dispersal throughout North America during this period (*66*). It is possible bison connectivity entering the Holocene may have been fostered by an expansion of C_4_ grasses (*67*) and changing behavior associated with body size diminution (*58*, *68*). Today, however, herd origin dictates bison population structure, in sharp contrast to the majority of bison history since the Last Glacial Maximum.

The population connectivity observed over the Holocene renders the apparent Mid-Holocene emergence of wood bison particularly notable; however, the precise timing, causes, and geographic origin of this emergence remain unclear. The Mid-Holocene represented a period of increasing aridity across portions of the North American plains (*69*), termed the Holocene Climate Optimum, which likely had adverse effects on bison population sizes (*70*, *71*) and also may have corresponded with a contraction of suitable bison habitat (*49*). After the termination of the Holocene Climate Optimum, woodland expanded at plains-boreal forest interface as the climate became cooler and wetter (*72–76*). This coincides with the date of our earliest wood bison, perhaps implicating such climatic variability as a factor driving subspecies divergence.

Human and bison populations have also been inextricably linked for thousands of years (*77*), and it is possible that shifting hunting pressures also played a role in mediating the trajectories of wood and plains populations. As morphological distinctions between the subspecies are imprecise (*10*), subspecies identification of fossil remains is challenging without genetic data.

Deeper sampling, particularly at the plains-boreal forest transition where wood bison occur in our data, would provide additional insight into the evolutionary history of wood bison.

All modern wood bison herds exhibit evidence of recent hybridization with plains bison, another legacy of recent bison management. The plains bison ancestry present in wood bison is specifically related to modern Elk Island plains herds, which originated from the same herd that was translocated to Wood Buffalo in the 1920s, pinpointing this event as facilitating subspecies admixture. Plains ancestry was variable across modern wood bison populations and subpopulations, with areas in close geographic proximity to the historical plains bison translocation center having higher amounts of this ancestry. The modern Elk Island and Mackenzie wood bison herds have the lowest plains bison ancestry, likely because they were populated with bison originating from remote areas of Wood Buffalo which were farther from the plains bison introduction site (*10*, *43*). The gradient of plains bison ancestry found in wood bison populations can be used to guide strategic, facilitated gene flow between herds striving to conserve wood bison ancestry. This variation may also offer insight as to whether there are any functional effects of, or selection for or against, plains bison ancestry in wood bison (*78*).

Despite their recent divergence and human-mediated gene flow, we found that wood bison today are highly distinct from plains bison, with the greatest genetic differentiation among modern bison populations — and over all post-Last Glacial Maximum North American bison — identified between subspecies.

Cattle introgression was believed to have been a major feature of recent bison population history as a result of intentional hybridization efforts by private herd managers (*20*). Although any potential ecological or biological effects of cattle ancestry are unknown, this has shaped both management policy (*26*) and public perception of bison as native North American wildlife (*79*). In addition to the limitations inherent in previously used genetic techniques (*19*), precisely detecting cattle ancestry has been challenging without access to genomes from bison that existed before cattle were introduced to North America. Using ancient bison as a baseline, we found that while there is evidence of individuals with cattle ancestry in several of the herds examined, its prevalence has been overstated. Moreover, there is substantial variation in the presence and frequency of cattle ancestry in our sampling, both within and across herds. Despite limited and intentionally biased sampling for some conservation herds, the majority of herds we examined contained at least one individual without cattle ancestry. Some groups, including Yellowstone-origin bison and all wood bison, had no evidence of cattle introgression. When present, cattle ancestry was found at low proportions and almost exclusively existed in a heterozygous state, implying that such ancestry may eventually be lost through recombination and drift in the absence of additional hybridization or selection.

Though bison are no longer facing immediate extinction risk, they remain heavily managed and are commonly confined behind fences. The majority of North American bison are raised for commercial production, with only a small proportion of individuals in herds whose purpose is to preserve the species’ diversity as native North American wildlife (*24*). While species such as elk [*Cervus canadensis*; (*80*)], bighorn sheep [*Ovis canadensis*; (*81*)], and pronghorn [*Antilocapra americana*; (*82*)] have more thoroughly returned to their role as wildlife, there is much work remaining to restore bison as important members of the Great Plains ecosystem (*83*, *84*) with opportunities to influence nutrient cycling, fire regimes, and overall biodiversity (*85–89*). Bison also play a vital cultural role in North America, particularly for Indigenous plains groups, and these human relationships with bison must be similarly restored (*90–94*). Paleogenomics offers another possible avenue for contextualizing these deep socio-ecological connections.

Best management practices for bison are difficult to identify without the broader lens into population history offered by ancient genomes, which capture patterns of diversity that are no longer present in modern herds. Characterizing such historical diversity allows managers to better understand the impacts of bison conservation and to optimize future restoration strategies (*27*). Some anticipated effects of the population collapse and recovery of bison, including the loss of genetic diversity and the ubiquity of cattle ancestry (*20*), may not be as dramatic as previously believed, whereas the increase in genetic differentiation across herds appears to be the most prominent signal separating pre-bottleneck bison from present-day populations. Given the intensity of human management during the recovery period, such differentiation is likely to have been driven by the isolation of small populations following the establishment of conservation herds, rather than by local adaptation or preexisting geographic structure. Heightened by the fragmented landscape bison have occupied along with the practical challenges of moving bison between geographically disparate herds, restoring genetic connectivity through management as a metapopulation would ensure the maintenance of genetic diversity (*8*), mitigate the fixation of cattle ancestry within herds, and more closely mirror how bison existed before their near-extinction. Metapopulation management would also reduce population structure across herds and could be crucial for facilitating the adaptation of bison populations to a wide variety of environments, including potential impacts of climate change (*95*).

This work offers a systematic overview of past and present bison diversity across North America since the Last Glacial Maximum to address key questions that have plagued managers as they attempt to stave off the loss of diversity in a highly fragmented modern continental bison population, while considering the impacts of inter- and subspecific introgression. Furthermore, we demonstrate the value of population-scale ancient DNA sequencing of wild species in disentangling the effects of recent anthropogenic pressures to support restoration efforts and effective management into the future.

## Materials and methods summary

For 115 ancient bison samples, we extracted DNA from ∼50mg of bone powder following a bleach pretreatment (*96*) using a protocol optimized for degraded DNA (*97*). We then made single-stranded libraries (*98*, *99*) for whole-genome sequencing. We also performed whole-genome sequencing on 45 modern bison samples, using libraries constructed from DNA extracted from hair or tissue. This dataset was combined with 52 previously generated bison genomes. Remnant adapters were trimmed and, for ancient samples, overlapping read pairs were merged. Reads were then mapped to the ARS-UCD1.2 cattle genome with the Btau5.0.1 Y chromosome added. For some introgression analyses, data was also mapped to the ARS-UCSC1.0 bison genome. To ascertain a set of single nucleotide polymorphisms encompassing bison diversity, we called variants in seven high coverage bison genomes: a modern wisent (*38*), a modern plains bison from Yellowstone National Park (*34*), a modern wood bison from Elk Island (*33*), four ancient Beringian steppe bison. This yielded a total of 3.6 million mappability-filtered transversions. Pseudohaploid genotypes were called for all samples at this set of sites using pileupCaller (https://github.com/stschiff/sequenceTools). We visualized genetic relationships using a t-SNE analysis (*100*), which we performed using Rt-SNE (https://github.com/jkrijthe/Rt-SNE) on a matrix of all possible *f4*-statistics relating individual bison, calculated with custom software (https://github.com/jooppenh/fstats/fstats.py). We also investigated population differentiation through time by measuring F_st_ between pairs of populations, and examined modern herd structure clustering using pairwise outgroup *f3-*statistics. Recent bison demographic histories were reconstructed using GONE2 (*57*). Conditional heterozygosity was calculated through pseudodiploid sampling to compare levels of genetic diversity across ancient and modern bison individuals. We used *D*-statistics and *f4*-ratios to test for and quantify plains bison ancestry in modern wood bison herds. To identify segments of cattle ancestry in modern bison genomes, we used a Hidden Markov model-based local ancestry inference approach (*64*). We constructed two ancestry source panels: one from four high-coverage steppe bison genomes and another using 20 modern cattle genomes from four breeds. We identified sites with at least 90% frequency difference between the two panels, and filtered these to be fixed in a large panel of cattle genomes, confidently call heterozygous ancestry in a F1 bison-cattle hybrid, and not have any reads carrying the cattle allele in any bison older than 1000 years. This left 5.36 million ancestry informative markers.

## Acknowledgements

We would like to thank the following people for facilitating sampling: B. McCann, National Park Service, for samples from Theodore Roosevelt National Park; M. Dooley and C. Ros for samples from Drumheller, AB; G. Zazula for samples from Yukon Territory; J. Driver for samples from British Columbia; I. Kirillova for Siberian samples. We are grateful to Q. Sakihara for assistance with data generation.

## Author contributions

Conceptualization: JO and BS with input from GAW, TKS, and GPA

Data curation: JO, SF, CNJ, DMD, and GAW

Formal analysis: JO

Funding acquisition: BS, GAW, TKS, and GPA

Investigation: JO, JDK, MC-J, SS, WES, HCC, SF, CG, LDB, and JS

Resources: AP, KPC, LCT, SRP, GM, KF, MAE, MCB, DRWB, JZM, JWI, RJL, TN, TP, CW, CML, KH, WO, PS, CNJ, DMD, and LCJ

Supervision: BS, GAW, LCJ, and REG

Visualization: JO

Writing – original draft: JO, LCJ, GAW, and BS

Writing – review & editing: all authors

## Data and materials availability

Sequencing data generated in this study is available through the NCBI Short Read Archive under BioProject accession no. PRJNA1374439.

## List of Supplementary Materials

Materials and Methods

Figs. S1 to S21

Tables S1 to S8

References *(101-153)*

