## Supplementary Materials for "Paleogenomic insight into the collapse, recovery, and management of American bison"

### Materials and Methods

#### Sampling

Our sampling was aimed at broadly surveying past and present North American bison diversity throughout their range since the Last Glacial Maximum. To contextualize their recent population decline, this sampling effort had a particular emphasis on ancient bison from the last few thousand years and modern bison from unsampled herds. We defined "modern" bison as those which lived within the last ~100 years, following their recent population collapse, while those that existed before and during the collapse were considered "ancient". These two categories also generally separate bison by DNA preservation, though there are samples from historical bison that lived after the bottleneck that also present characteristics typical of ancient DNA (short fragment lengths, low levels of endogenous DNA, and cytosine deamination). Metadata for newly sequenced ancient and modern samples is presented in table S1.

The locations of modern bison herds featured in this study are presented in fig. S1. The following abbreviations are used for these herds at times throughout, largely following Hartway *et al*. (*8*):

- **Plains bison**: CCSP = Caprock Canyons State Park; SCI = Santa Catalina Island; NER = National Elk Refuge/Grand Teton National Park; FTN = Fort Niobrara National Wildlife Refuge; VPR = Vermejo Park Ranch; WICA = Wind Cave National Park; WMW = Wichita Mountain Wildlife Refuge; YELL = Yellowstone National Park; ELKp = Elk Island National Park plains bison.
- **Wood bison**: ELKw = Elk Island National Park wood bison; MBS = Mackenzie Bison Sanctuary; RLBH = Ronald Lake Bison Herd; WAB = Wabasca; WBNP = Wood Buffalo National Park; For some analyses, the WBNP herd is split into geographic subpopulations: GR = Garden River; SRL = Slave River Lowlands; PAD = Delta; NY = Nyarling; HC = Hay Camp.

We sampled several individuals that were of Yellowstone origin but are managed in other herds, including the Fort Peck bison herd and the Laramie Foothills herd, via the Rocky Mountain Arsenal National Wildlife Refuge. These individuals were considered part of the YELL herd for herd-level analyses.

<u>Bison of the Bighorn Basin Project</u>

In 2020, the Bison of the Bighorn Basin Project was launched as a collaborative research initiative involving local community members who donated their privately owned bison skulls for analysis by a team of researchers (https://meeteetsemuseums.org/tag/bison-of-the-bighorn-basin/). The project goal was to develop a detailed dataset of bison from the geographic Bighorn Basin to better understand their ecology prior to the 19th-century population bottleneck. In total, 114 bison were recorded, with 24 sampled for radiocarbon dating and stable carbon and nitrogen isotope analysis based on completeness (Phillips et. al., in press). Eleven of the radiocarbon-dated crania were selected for the present study (table S1).

**Sequencing ancient and modern bison genomes**

We removed the surface from a small portion of each bison bone sample with a Dremel 400 rotary drill (Dremel, Mt. Prospect, IL), then cut off a ~100-500mg subsample. We crushed the subsamples using a Retsch MM 400 mixer mill (Retsch GmbH, Haan, Germany) and then extracted DNA from ~50mg of the resulting bone powder using the silica column-based buffer D option from Rohland *et al.* (*97*), following a 0.5% bleach pretreatment (*96*). From the DNA extracts, we prepared dual-indexed Illumina sequencing libraries (*101*), either manually or on an Agilent Technologies Bravo NGS Workstation (Agilent, Santa Clara, CA), using a single-stranded protocol that was designed for the conversion of degraded DNA (*98*), with modifications from Nguyen *et al*., 2023 (*99*). Library preparation was performed without enzymatic repair of ancient DNA damage (*102*) in order to maximize the complexity of recovered DNA. All pre-PCR ancient DNA molecular work was carried out in a dedicated PCR-free clean room at the UCSC Paleogenomics Lab.

We extracted DNA from modern bison hair or tissue samples using the Qiagen DNeasy blood and tissue kit (Qiagen, Venlo, The Netherlands). Extracted DNA was then converted into Illumina sequencing libraries using the NEBNext Ultra II FS kit (NEB, Ipswich, MA), with 7- or 8-basepair dual-index barcodes (*101*). In the case of several hair samples that yielded short DNA fragments, we instead used the same single-stranded ancient DNA library preparation method that was used for ancient samples.

For both ancient and modern molecular work, samples were grouped into batches of 4-24 for extraction and library preparation. Negative controls were included in each extraction and library preparation batch.

We quantified ancient and modern libraries using the Qubit 1x HS kit and assessed fragment length distribution with either the 5300 Fragment Analyzer or the Tapestation 2200 (Agilent, Santa Clara, CA). The endogenous content and complexity of each library was estimated by performing low-depth sequencing (~1M reads) on an Illumina NextSeq 550 or NextSeq 2000 at the UCSC Paleogenomics Lab. Samples with adequate preservation were selected for whole genome deep sequencing using a NovaSeq 6000 or NovaSeq X at either the University of California, San Francisco Center for Advanced Technology or the Duke University Genomic and Computational Biology sequencing core.

### Radiocarbon dating

For 75 bison that had not been previously dated, ~100-1000mg subsamples of bone or bone powder were dated at the UCI Keck AMS Laboratory, using the remaining material from the subsample used for DNA when possible. Two samples failed to yield sufficient collagen for radiocarbon dating, leaving 73 samples from which we obtained a date. Radiocarbon dates were calibrated with the IntCal20 Northern Hemisphere atmospheric curve (*103*) using OxCal 4.4 (*104*). Isotopic results are presented in table S3.

### Incorporating publicly available bison genomes

We combined our ancient and modern bison dataset with a set of 52 publicly available modern American bison (*Bison bison bison* and *Bison bison athabascae)* genomes *(20, 33–39)*, summarized in table S2. We also included modern and historic wisent [*Bison bonasus; (38, 39, 105–107)*], and other bovid species, including taurine (*Bos taurus*) and indicine (*Bos indicus*) cattle from six breeds (*36*, *108–110*), yak [*Bos grunniens*; (*111*)], gaur [*Bos gaurus;* (*36*)], and water buffalo [*Bubalis bubalis; (112)*].

### Data processing and bioinformatics

We adopted separate bioinformatics processing pipelines for ancient and modern samples to address issues associated with short fragment lengths and chemical damage present in degraded specimens. For ancient samples, we trimmed remnant adapter sequences and merged overlapping read pairs using SeqPrep2 (https://github.com/jeizenga/SeqPrep2), requiring a minimum length of 35 base pairs. Unmerged reads were discarded. We then mapped them using bwa v0.7.17-r1188 aln (*113*) with parameters optimized for ancient DNA ( -n 0.01 –o 2 –l 16500). For modern samples, we trimmed adapters using Trimommatic v0.39 (*114*) and mapped reads using bwa MEM (*115*) with default parameters. In both cases, we mapped reads to the ARS-UCD1.2 cattle genome (*109*), with the Btau5.0.1 (GCF_000003205.7) Y chromosome added. We then removed duplicate molecules using Picard MarkDuplicates (https://broadinstitute.github.io/picard/). For ancient samples, we measured the rate of deamination-induced C->T mismatches at the terminal ends of molecules using mapdamage2 v2.0.6 (*116*) to confirm the authenticity of the DNA.

To obtain a set of single nucleotide polymorphisms (SNPs) for analysis, we called variants in seven high (≥17-fold) coverage bison genomes: a modern wisent (*38*), a modern plains bison from Yellowstone National Park (*34*), a modern wood bison from Elk Island National Park (*33*), two ancient Siberian steppe bison (*Bison priscus*; ~46 and 10 thousand years ago; ka), and two ancient steppe bison from the Yukon (~50 and 23ka). For the four ancient steppe bison, we used both merged and properly-paired reads for genotyping. We called genotypes separately for each individual, using GATK v4.1.8.1 HaplotypeCaller (*117*) for modern samples and snpAD (*118*), an ancient DNA-aware variant caller, for the steppe bison. As snpAD does not perform local realignment, we used GATK RealignerTargetCreator and IndelRealigner (v3.8-1-0-gf15c1c3ef) to realign areas around indels for the four steppe bison prior to variant calling.

We then filtered each set of variants with BCFtools v1.8 (*119*) using the +setGT plugin for a minimum genotype quality of 30 and minimum and maximum depths of 1/3rd and twice the mean coverage on a per-sample basis. To obtain a final set of SNPs, we took all heterozygous variants called for each individual, as discovering variants present within the two chromosomes of a single individual provides an unbiased sampling of the population allelic spectrum (*120*). We then restricted to transversion variants, which cannot be caused by ancient DNA-associated deamination, and filtered for mappability using SNPable with a *k*-mer length of 35 and stringency of 0.5 (gen_mask -l 35 -r 0.5; https://lh3lh3.users.sourceforge.net/snpable.shtml). This yielded a final set of 3,569,680 filtered transversions across among these 7 ancient and modern bison. We called pseudohaploid genotypes on these variants for all samples using SAMtools v1.9 (*121*) mpileup and pileupCaller (https://github.com/stschiff/sequenceTools), requiring a minimum base quality of 30 (-Q 30) and minimum mapping quality of 25 (-q 25), with BAQ disabled (-B).

**Pruning relatives**

READv2 (*122*) was used to identify close relatives within modern bison herds. READv2 detects relatives using pairwise mismatch rates in pseudohaploid genotype data, with a baseline established from unrelated individuals from the same population. Relative detection was performed in wood bison and plains bison separately. Pseudohaploid EIGENSTRAT genotype files were converted to PLINK format with convertf (https://github.com/DReichLab/AdmixTools), subset to only modern samples, and split by subspecies. Genotypes were filtered using PLINK v1.9.0-b.8 (*123*) to require a minor allele frequency of 5% and a maximum per-variant missingness of 33%. READv2 was then run using default parameters.

As modern bison display stronger population structure between herds (described in 'Genetic structure and population connectivity' below), mismatch rates within and and across herds may differ even for unrelated individuals. Given low sample sizes and unknown patterns of relatedness for some herds, using herd-specific baselines was not always feasible. Therefore, we compared relative detection using as a normalization value either the median mismatch rate across all modern bison or the median mismatch rate for 18 roughly contemporaneous Late Holocene Interior bison (295-102 calibrated years before present).

For three herds with elevated levels of inbreeding (described in 'Genetic diversity and inbreeding' below), long runs of homozygosity (ROH) could reduce mismatch rates between individuals within these herds and result in unrelated individuals being identified as close relatives. We therefore performed relative detection separately within these three herds: the

Caprock Canyons State Park plains bison herd (CCSP; n=6) and the Wabasca (WAB; n=6) and Ronald Lake (RLBH; n=6) wood bison herds. For these herds, we ran READv2 on a per-herd basis and used the maximum mismatch rate among individuals for normalization (-n max). We also compared this within-herd relative detection using READv2 to that obtained using KIN (*124*), which estimates relatedness by modeling segments in the genome that are identical by descent and identifies runs of homozygosity. We ran KIN within each herd without contamination correction (-cnt 0) across all autosomes (-N 29).

Overall 13 pairs of first-degree relatives were detected. This approach correctly identified two known parent-offspring pairs from the Fort Niobrara National Wildlife Refuge herd: 0618 as the offspring of 1320 and 0419 as the offspring of 1535 (table S1). First-degree relative detection was consistent using either the mismatch rate baseline derived from modern bison or that from recent ancient plains bison, with the only difference being that one pair of Elk Island wood bison (W2 and W3) was identified as a first-degree relative pair using the modern wood bison baseline but was not with the ancient plains bison baseline. For the WAB and RLBH herds, both READv2 with the within-herd max mismatch rate normalization and KIN detected no first-degree relative pairs (while 9 and 2 pairs, respectively, were detected using the overall modern wood bison median mismatch rate with READv2). The estimates for the CCSP herd between READv2 with the max mismatch rate and KIN were largely consistent, with 9 and 10 pairs of first-degree relatives detected, respectively, of which 8 overlapped.

One individual from pairs of first-degree relatives were removed from the dataset for downstream dimensionality reduction and allele-sharing analyses. Relative pruning prioritized removing individuals with higher numbers of relatives across pairs or, if equal, individuals with lower sequencing coverage. First-degree relative pairs detected across modern bison are summarized in table S4.

We also ran READv2 in ancient bison within major temporal and geographic groups of samples, using default parameters. We identified one pair of samples from Fort Walsh as originating from the same individual or from identical twins: 448N1C99-1 and 448N1B1-1. We retained 448N1B1-1, which had greater sequencing coverage, for downstream analyses.

**Genetic structure and population connectivity**

We visualized genetic relationships across all sampled bison by performing principal component analyses (PCA) on a matrix of all possible *f4*- or *f3*-statistics (*40*) of the form *f4(X, A; B, C)* and *f3(X; A, B)*, where X, A, B, and C are different individual bison (Fig. 1B). This approach allowed for observing relationships between bison without relying on projecting low-coverage ancient samples onto axes of modern genetic variation (*125*), as is common in ancient DNA studies (*126*), since we expected modern variation may incompletely reflect the diversity of past bison populations (*127*). PCA was performed using the R function prcomp with scale=TRUE. We also used the resulting principal components as input for t-SNE (*100*) and UMAP (*128*) analyses, which allowed for viewing in two dimensions patterns that were visible in higher principal components. We performed the t-SNE and UMAP in R with the Rt-SNE (https://github.com/jkrijthe/Rt-SNE) and umap (https://github.com/tkonopka/umap) packages. Due to the number of statistics involved, *f*-statistics for dimensionality reduction analyses were calculated with custom software (https://github.com/jooppenh/fstats.py), which uses the LLVM-based numba just-in-time Python compiler (*129*) and NumPy arrays (*130*) to calculate large numbers of *f*-statistics efficiently. For the *f3*-statistic-based analyses, the seven individuals involved in variant ascertainment were omitted, as these individuals displayed slightly higher *f3*-values as the target population—essentially having longer tip branch lengths due to this ascertainment scheme. Because *f4*-statistics measure internal branches within trees, they are potentially less sensitive to differences in tip branch length and so these individuals were retained for the *f4*-based analyses (*131*, *132*). Visualizations of these different dimensionality reduction approaches using either *f4*- or *f3*-statistics as a basis are presented in fig. S2.

We also performed a similar *f4*-statistics-based PCA using only ancient bison samples in order to assess genetic structuring since the Last Glacial Maximum in a way that was not complicated by recent bison management (Fig. 2A). We observed that ancient bison were primarily grouped by time in the PCA, suggesting that there was largely temporal rather than geographic stratification in our sampling of ancient bison genomes, reflecting a highly connected population over much of North America during this period. We then assessed this relationship by calculating the Pearson's correlation coefficient between the position of each sample on PC1 and the age of the sample.

Two groups stood out as possible exceptions to this temporal structuring: remnant northern steppe bison populations and the wood bison subspecies, although individuals within these groups also followed the overall trend relating sample age and genetic structure. These remnant northern bison were shifted upward along the first t-SNE axis, perhaps because they had some ancestry component not seen in other North American bison and which was retained in these individuals into the Holocene. One possibility was that this ancestry component was associated with Siberian bison, which could have been spread by movement across the Bering Land Bridge during the Ice Age (*51*, *54*, *133*). We tested for such Siberian-related ancestry using the statistic *f4(Interior Early Holocene; northern; Siberia 46ka, wisent)*, which tests whether northern bison have excess affinity to a pre-LGM Siberian steppe bison relative to ~10ka bison from Wyoming. We found that all northern bison have significant ($Z>3$) allele sharing with this Siberian bison, suggesting that Siberian-related ancestry explains the position of these individuals in the t-SNE analyses (fig. S4).

Wood bison were the other bison group with distinct genetic structure in dimensionality reduction analyses. All modern wood bison formed a separate cluster relative to plains bison, indicating that the two bison subspecies correspond to discrete genetic units. Fifteen ancient bison were shifted toward this wood bison cluster, compared to other Holocene Interior bison (Fig. 1B). This shift suggested that these bison had wood bison ancestry, which we confirmed using allele sharing statistics testing for affinity between these bison and modern wood bison (fig. S3). The first ancient bison which grouped with modern wood bison had a date of 2910 calBP. This date was perhaps more recent than expected given the subspecies designations of the two groups, and supports a Mid-Holocene origin for wood bison. After their divergence, it is possible that there was recurrent contact between subspecies in areas where their ranges overlapped in central Alberta and British Columbia (*43*). To examine the potential for past gene flow between ancient wood and plains bison, we tested whether all ancient wood bison formed a clade relative to modern Elk Island plains bison using *D*-statistics of the form *D(historical WBNP, ancient wood; ELKp, wisent)* for all ancient wood bison. These statistics were consistent with 0 for all ancient wood bison (fig. S11), suggesting that there was no recurrent contact between subspecies following their divergence. We also found no wood bison in the more southerly areas of their previously estimated historic distribution (*43*), such as central Alberta around Edmonton (Fig. 4A), and so their range may have been restricted to the margins of the plains-boreal forest interface. We did, however, observe wood bison in Banff and Winnipeg, which could indicate wood bison were more widely spread at this grassland-forest transition zone than has been assumed (Fig. 1A).

As a point of comparison for the allele sharing-based dimensionality reduction analyses, we also used PCAngsd v1.36.4 (*134*) to perform PCA using genotype likelihoods.This approach also allows for including ancient individuals in calculating principal components, and does not rely on ascertainment in a set of high-coverage genomes, as was used for generating the pseudohaploid dataset. For all individuals with ≥0.5-fold coverage, we calculated genotype likelihoods using ANGSD v0.941-22-gc877e7f (*135*), with the parameters -GL 1 -doMajorMinor 1 -doMaf 2 -SNP_pval 1e-6 -minMapQ 25 -minQ 30 -minInd 26 -minMaf 0.05 -doGlf 2 -remove_bads 1 -uniqueOnly 1 -rmTrans 1, restricting to mappable regions (from the SNPable mappability filter) using the -sites flag. This resulted in a total of 4,535,642 transversion variants with minor allele frequency ≥ 0.05. PCAngsd was run with default parameters.

The PCAngsd analysis largely agreed with those based on *f*-statistics, in which there was a strong temporal gradient and distinct groups representing remnant northern steppe bison and wood bison along with Interior and plains bison (fig. S5). PC1 separated ancient and modern individuals, which likely reflects to some degree recent drift which has occurred in modern populations. However, PC1 position also seemed to correspond to sample quality, such that modern bison with shorter fragment sizes (data for which was in most cases generated using ancient DNA-specific molecular methods optimized for degraded DNA) were shifted towards ancient samples on PC1. Therefore, we concluded that PCAngsd is more sensitive to technical biases than the *f*-statistics-based dimensionality reduction analyses.

Wyoming time transect

To examine how North American bison population relationships changed over the Holocene, we assembled a time transect of 16 bison from 5 archaeological sites across Wyoming. These sites included: Horner (n=5; ~10.5ka), Finley (n=3; ~10.2ka), Carter Kerr-McGee (n=3; ~10ka), Hawken (n=3; ~7ka), and Scoggin (n=2; ~4.1ka), along with 11 bison from Wyoming's Bighorn Basin dating between 690-100ya (*55*, *136–138*). These bison samples span essentially the entirety of the Holocene (Fig. 2B), and so comparing how these samples were related to others in our dataset provided a lens for examining population changes over this period. We did this by measuring relative allele sharing between pairs of sites adjacent in age to all other sampled bison genomes X, using statistics of the form *f4(wisent, bison X; younger WY site, older WY site)*. *f4*-statistics were calculated using admixtools2 (*139*).

Given connectivity between Wyoming and other bison populations, we would expect that individuals which postdate both sites would share more alleles with the younger site than the older one. These comparisons are presented in Fig. 2B, excluding Carter Kerr-McGee, both to facilitate visualization and because it had a wider range of radiocarbon dates between samples and one sample which failed dating.

This pattern of connectivity, in which younger bison shared more alleles with bison from the younger site than the older site, was found throughout the Holocene. Such allele sharing is greater with larger temporal gaps between sites, but is also seemingly present at a fine scale when comparing sites close in time (such as Horner I, ~10.5ka, and Finley, ~10.2ka, in which younger bison have greater allele sharing with Finley). This allele sharing agrees with the temporal structuring in the t-SNE analysis in demonstrating a largely panmictic population of bison across the Interior throughout the majority of the Holocene. In the absence of such connectivity, it would be expected that geographically distant bison would be equally related to these Wyoming bison across all time points. Instead, this temporal gradient demonstrates continual genetic linkages between Wyoming bison and those across the Interior. However, it is possible that such connectivity does not necessitate extensive long-range mobility at an individual level, and there were likely differences in mobility across past bison populations, including the Wyoming bison examined here, given variation in forage and water availability.

Genetic vs. geographic distance through time

To compare differences in how genetic and geographic differentiation corresponded over time in North American bison since the LGM, we divided the last ~13,000 years into four distinct temporal segments: the Pleistocene-Holocene transition (~13-10ka), Mid-Holocene (~7-3ka), Late Holocene (~3-0.1ka), and modern. Individuals were grouped into populations based on temporal and geographic proximity.

The following populations were formed for each time segment:

- **Pleistocene-Holocene transition**: Yukon_13k (n=7), CloverBar_AB_13k (n=5), Farr_SK_11k (n=2), Horner_WY_11k (n=5), Finley_WY_10k (n=3), CKM_WY_10k (n=3)
- **Mid-Holocene**: Hawken_WY_7k (n=3), Central_AB_4k (n=6), Scoggin_WY_4k (n=2), TRNP_ND_3k (n=2), Yukon_3k (n=3)
- **Late Holocene**: Central_AB_500 (n=3), MT_SK_100 (n=3), RidingMtn_MB_800 (n=3), Rockies_MT_CO_700 (n=3), wood_AB_200 (n=5), BighornBasin_WY_200 (n=11), FtWalsh_SK_2k (n=2), kā-kī-māmawēpihk_SK_300 (n=5), Drumheller_AB_300 (n=3), WBNP_historic (n=2)
- **Modern**: YELL (n=15), ELKp (n=4), FTN (n=2), WMW (n=2), SCI (n=2), CCSP (n=2), VPR (n=2), WICA (n=3), WBNP (n=9), MBS (n=3), ELKw (n=8), RLBH (n=6), WAB (n=6)

Within each time point, we compared geographic and genetic differentiation for all population combinations (Fig. 2C). For geographic distance, we calculated the Haversine distance in kilometers between the average geographic position of individuals within each population using distm() from the geosphere R package (https://github.com/rspatial/geosphere). As a measure of genetic differentiation, we computed $F_{st}$ using admixtools2 (*139*) using the fst() function with the parameters maxmiss=1 and auto_only=F.

Modern herd population relationships

We used pairwise outgroup *f3*-statistics of the form *f3(wisent; herd 1, herd 2)* to understand population structure among modern wood and plains bison herds (Fig. 3B). The Wood Buffalo herd was split into subpopulations, and all individuals from production herds were grouped into a single population (n=13). As these statistics measure shared drift between pairs of herds, they are potentially less sensitive to drift associated with individual bottlenecks present in the foundational histories of many herds. Outgroup *f3*-statistics were calculated from genotype files using admixtools2 (*139*) with the f3() function, setting auto_only=F and outgroupmode=T. Pairwise outgroup *f3* values are presented in Fig. 3B, along with a dendrogram generated using hierarchical clustering with the hclust() R function using a dissimilarity matrix of (1-*f3*) for all pairwise values.

**Genetic diversity and inbreeding**

To obtain a measure of individual-level genetic diversity, we performed pseudodiploid genotype calling by sampling two reads for each individual across the set of 3.6M ascertained bison transversions using pileupCaller (--randomDiploid), following the same filtering criteria described in the 'Data processing and bioinformatics' section. We then estimated conditional heterozygosity as the rate of heterozygosity among the pseudodiploid calls, requiring a minimum of 10,000 SNPs to be covered by at least two reads for each sample (Fig. 3C).

We also estimated runs of homozygosity (ROH) in modern bison with sufficient (~5-fold) coverage using ROHan (*140*). We ran ROHan with the option --rohmu 5e-4 for each individual, within autosomal regions in the SNPable mappability filter using the --bed flag and with a window size of 1 megabase. We then calculated the Pearson's correlation coefficient between the proportion of the genome that is contained in ROH and the estimated conditional heterozygosity rate, and found that differences in ROH content are highly correlated with variation in estimated heterozygosity (fig. S7; $r_{\text{heterozygosity,\% ROH}} = -0.78$, $P = 1.01 \times 10^{-14}$). We also compared the pseudodiploid conditional heterozygosity rate and the theta values estimated with ROHan (including ROH in estimated theta), and found that they largely agree, although with some outliers (fig. S8). We then estimated the per-sample error rate, using as a proxy the rate of non-biallelic bases appearing in reads at the ascertained bison transversions (i.e. the rate of reads carrying neither of the two ascertained alleles at these sites), and observed a relationship between the ROHan-estimated heterozygosity (excluding regions in ROH) and the error rate, with the outliers found in comparing the ROHan and pseudodiploid heterozygosity estimates largely having the highest estimated error rates (fig. S9). Pseudodiploid conditional heterozygosity estimates did not display this same relationship (fig. S10). We therefore interpret that ROHan may be more sensitive to varying levels of sequencing or mapping error across samples in estimating heterozygosity.

**Demographic reconstruction**

We reconstructed the demographic trajectories of modern and recent wood and plains bison over the last ~100 generations using GONE2 [Fig. 3A; (*57*)]. We used the modern herds with the largest sample sizes for demographic reconstruction of each subspecies: Yellowstone National Park origin bison (n=15) for plains bison and the Wood Buffalo National Park herd (n=9) for wood bison. Demographic trajectories inferred using only these modern herds were compared to estimates derived from recent ancient populations, using 18 Interior bison with calibrated radiocarbon dates of less than 300ya for plains bison and 11 ancient wood bison <1ka, requiring a minimum of 0.3-fold coverage for each sample (median of 1.73-fold and 2.31-fold coverage for the ancient plains and wood bison, respectively). We subset genotype files on a per-group basis using PLINK and removed sites missing across all individuals within each population.

We ran GONE2 separately for each herd with the following parameters: -g 3 -r 1.0 -b 0.001 -e, which uses pseudohaploid data and assumes a flat recombination rate. In plotting, we shifted the curves estimated for the ancient populations back to align their curves with those of modern populations. This resulted in a shift of 15 generations for the 300ya interior bison and a shift of 35 generations for ancient wood bison. These shifts agree closely with the mean calibrated radiocarbon dates for each group, assuming a generation time of ~10 years (*58*): 179 calBP for plains bison and 327 calBP for wood bison.

**Subspecies introgression**

Due to historical translocations of plains bison into Wood Buffalo National Park (WBNP) (*15*), modern wood bison may have plains bison ancestry not seen in past wood bison populations. We sought to evaluate the impact of these translocations on modern wood bison ancestry using our sampling of wood bison which existed before these events took place. We used D-statistics of the form *D(historical WBNP, modern wood bison; Elk Island plains, wisent)* to detect excess plains ancestry in modern wood bison relative to two historical wood bison samples from WBNP dating to ~175 years before present and collected before the plains bison translocation into WBNP (Fig. 4B). To quantify the amount of plains introgression in modern wood bison herds, we used an *f4*-ratio test of the form *f4(ancient wood bison, Siberia Pleistocene; modern wood herd X, Elk Island plains)/f4(ancient wood bison, Siberia Pleistocene; historical WBNP, Elk Island plains)* for each modern wood bison herd, in which we grouped all ancient wood bison samples into the ‘ancient wood bison’ population, except for the two historical WBNP individuals (historical WBNP). This ratio measures the amount of plains ancestry present in each modern wood bison herd, using the historical WBNP samples as a baseline for wood bison ancestry and the Elk Island plains herd as a potential source, as Elk Island plains bison are derived from the same herd that was translocated to WBNP (Fig. 4C and Fig. 4D).

To confirm Elk Island as the source of introgression, we used D-statistics of the form *D(Elk Island plains, modern plains herd X; modern wood herd Y, wisent)*, which tests for excess affinity between Elk Island plains bison and modern wood bison herds relative to other modern plains herds, including the Yellowstone National Park, Wind Cave, Wichita Mountains, Fort Niobrara, Caprock Canyons, Catalina Island, and National Elk Refuge herds (Fig. 4E). We also excluded the possibility of this signal being driven instead by introgression from wood bison into Elk Island plains bison by testing for allele sharing between modern wood bison herds and other plains herds, using statistics of the form *D(historical WBNP, modern wood herd, modern plains herd, wisent)* for all non-Elk Island plains bison herds (fig. S12). All D- and *f4*-statistics statistics for evaluating plain bison ancestry were calculated using admixtools2 (*139*).

We used the qpWave (*63*) framework implemented in admixtools2 to estimate the minimum number of sources of plains bison ancestry across all wood bison herds. As possible plains bison sources (‘right’ populations), we used following minimal set of herds which captured the major groups seen in the outgroup *f3*-statistic clustering of modern bison:

- ELKp, YELL, WMW, WICA, FTN

For modern/historical wood bison (‘left’ populations), we used the following herds:

- RLBH, WBNP, ELKw, MBS, WAB, historical_WBNP (two ~175ya wood bison from WBNP grouped into a single population)

The qpWave results are presented in table S5. A model with no admixture is strongly rejected (p=6.12E-27), although a model with a single admixture event provides and adequate fit for the data (p=0.28), suggesting that one stream of gene flow between wood and plains bison is largely sufficient to explain plains bison ancestry in wood bison.

Next, we used qpAdm (*141*) to model the ancestry of modern wood bison herds. We used a rotating modeling approach (*63*, *142*) with the following set of source/reference populations:

- ELKp, YELL, WMW, WICA, FTN, historical_WBNP, wood_Late_Holocene

in which wood_Late_Holocene is a group of all other ancient wood bison (except the two ~175ya WBNP individuals used for the 'historical_WBNP' population), and the other populations represent the minimal set of plains bison ancestry sources that encompass major groups of plains bison.

We performed qpAdm modeling with each modern wood bison herd as the target using the rotate_model() and qpadm_multi() functions from admixtools2. Passing qpAdm models ($p \geq 0.01$ with admixture weights between 0 and 1) are presented in table S6. The qpAdm models agree with the estimates obtained from *f4*-ratios, in that there is a gradient of plains bison ancestry across modern wood bison. Both methods give consistent admixture proportion estimates, although slightly higher plains proportions are estimated using qpAdm. Modeling with qpAdm also indicates that Elk Island plains bison are the best source, as a two source model using historical_WBNP and ELKp fits for every herd. For the ELKw and MBS herds, YELL can be used in place of ELKp as a plains ancestry source, although this provides poorer fits ($0.01 < p < 0.05$). Finally, a wider range of three-source models also fit for the WAB herd, possibly suggesting they have an additional source of plains bison ancestry not seen in other wood bison, which could account for their higher proportions of plains ancestry.

Our qpAdm modeling consisted of both ancient and modern populations in the source/reference set, which creates some underlying *f4*-statistics which have a mix of ancient and modern individuals on both sides of the statistic, a known source of potential bias (*63*). To assess whether such bias may be influencing our results, we performed another round of qpAdm modeling of modern wood bison ancestry using the same rotating approach but with only ancient populations in the source/reference set:

- Yukon_Pleistocene, Central_AB_4k, wood_Late_Holocene, historical_WBNP Bighorn_WY_250

Passing models from this rotating, ancient-only qpAdm analysis are shown in table S7. Overall, the results are similar in which modern wood bison herds were best modeled as mixtures of a recent wood bison ancestry source (historical_WBNP), and a recent plains bison source, which consisted of 11 bison from Wyoming dating within the last 700 years (Bighorn_WY_250).

Another major question in wood bison population history was the origins of the Ronald Lake and Wabasca herds (*11*). We sought to elucidate the relationship of Wabasca and Ronald Lake to other modern wood bison herds using D-statistics of the form *D(ancient wood bison X, modern wood bison herd Y, Wabasca/Ronald Lake, wisent)*, which tests whether the Wabasca or Ronald Lake herds share more alleles with modern wood bison than they do with any ancient wood bison sample (fig. S13). If either the Wabasca or Ronald Lake herds represent an unsampled wood bison lineage, they would be expected to be either equally related to modern and ancient wood bison or to be closer to one of the ancient wood bison populations. Alternatively, if they are relatively recently derived from another wood bison herd, they would be expected to have a genetic association with modern herds, relative to ancient wood bison. However, since Wabasca or Ronald Lake may share more alleles with modern herds through recent plains bison introgression rather than population history, we also examined *D(ancient wood bison X, historical WBNP; Wabasca/Ronald Lake, wisent)*, as our pre-1928 bison from WBNP lack recent plains admixture but share a close genetic relationship with modern day wood bison herds. Both the Ronald Lake and Wabasca herds share more alleles with recent and historical wood bison, relative to ancient wood bison, suggesting that they are closely related to the modern WBNP herd and are not part of a separate wood bison lineage (fig. S13).

#### Cattle introgression

Due to deliberate attempts to hybridize bison and cattle during the early 20th century (*17*), it is possible that some, and perhaps even all (*20*), modern bison possess cattle ancestry. Our sampling of ancient bison which existed before cattle were introduced to North America provided a baseline for assessing the extent of recent interbreeding within and across modern bison genomes.

<u>D-statistic test for introgression using alignments to the bison reference genome</u>

We first used allele sharing statistics to test for evidence of cattle introgression in modern individual bison. As cattle are largely an outgroup to past bison populations, for other analyses we used data aligned to the cattle reference genome in order to avoid reference bias associated with ancient DNA (*143*, *144*) inflating similarity between ancient bison and the Yellowstone reference individual. However, such reference bias complicates detecting cattle ancestry, as artifactual similarity between ancient bison and the cattle reference could preclude the detection of small proportions of cattle ancestry in modern bison. Therefore, we also aligned a subset of ancient and modern bison to the ARS_UCSC1.0 bison reference genome (*34*), with the ARS-UCD1.2 cattle X chromosome and the bison mitochondrial genome (NC_012346.1) appended.

We ascertained variants in 10 medium and high (>6-fold) coverage modern bison genomes along with high coverage bovine genomes from a range of species and breeds: water buffalo (SAMN08640746), Hariana cattle (SAMEA5577149), Angus cattle (SAMN02843152), Charolais cattle (SAMN02843066), gaur (SAMN05558794), yak (SAMN03761424), wisent (SAMN08323725). Variants were called using GATK v4.1.8.1 HaplotypeCaller (*117*) in -ERC GVCF mode to generate per-sample GVCFs, which were then merged using CombinedGVCFs and genotyped using GenotypeGVCFs. We filtered genotypes on a per-sample basis to have GQ ≥ 30 and coverage between ⅓- and 3-fold the genome-wide average, with a minimum depth of 5, setting any genotype failing any of these criteria as missing. Using BCFtools view, we restricted this set of filtered variants to biallelic transversions which had called genotypes in at least ⅓ of the individuals (F_MISSING<0.66), yielding 23,187,604 SNPs. We then called pseudohaploid genotypes on all samples using mpileup/pileupCaller, with the same filtering criteria described in 'Data processing and bioinformatics', and used this pseudohaploid dataset for evaluating allele sharing between modern bison and cattle.

We calculated *f4*-statistics of the form *f4(modern bison, Horner_WY_10k; angus, gaur)*, which test for allele sharing between angus cattle and modern bison relative to a ~10.5ka bison from the Horner site in Wyoming to look for evidence of cattle introgression in 20 modern bison from 9 herds (Fig. 5A). Three bison had significant (Z > 3) allele sharing with cattle, providing evidence for cattle ancestry in some modern bison. However, the majority of bison exhibited no excess affinity with cattle. This was unexpected as it had been previously reported that all bison have some amount of cattle ancestry (*20*). Additionally, our allele sharing test failed to provide evidence of cattle introgression in several individuals that were identified as having cattle ancestry in Stroupe *et al.* (*20*), including the individual that was shown to have the most cattle ancestry in that study (HistW_1937/W13) .

To test the sensitivity of this analysis to the specific individuals used for calculating *f4*-statistics, we also calculated these statistics with different ancient bison, cattle breeds, and outgroup bovine species and obtained nearly identical results (fig. S14). To evaluate the impact of our specific variant ascertainment scheme, we also performed ascertainment within only bison (using the 10 medium and high coverage genomes) and again obtained highly similar results (fig. S15).

Local ancestry inference

Given the surprising lack of cattle ancestry in our initial introgression test, we next sought to survey the amount and location of cattle ancestry within all sampled bison genomes. For this, we used a Hidden Markov model-based local ancestry inference approach, ancestry_HMM (*64*, *145*), to assign diploid ancestry in individual genomes as originating from bison or cattle. As a source panel for bison, we used the 4 high coverage Late Pleistocene steppe bison from Siberia and the Yukon, and for cattle we used 20 moderate-to-high coverage genomes (>10-fold coverage) from 4 taurine cattle breeds [table S8; (*36*)]. We called variants using data aligned to the cattle reference genome in each panel separately, using snpAD to call individual-level all-sites VCFs for the steppe bison panel and GATK HaplotypeCaller to create gVCFs for the cattle, which were then merged with CombineGVCFs and genotyped with GenotypeGVCFs. We then filtered sites on a per-individual basis for a minimum genotype quality of 30 and coverage between ⅓- and 3-fold the genome-wide average, restricting to mappable regions. We then merged these two panels, requiring sites to be biallelic SNPs, be called in at least 2 ancient bison and 6 cattle, and have a frequency difference of at least 90% between the two source panels. This yielded 8,639,845 putative ancestry informative markers (AIMs). We filtered this set of AIMs by removing any which were variable among the 1000 Bull Genomes Project Run 9, a set of 488 high coverage cattle genomes (*65*), essentially requiring AIMs to be fixed in cattle. We also removed any which had <90% posterior probability of being heterozygous in “Buzz”, an F1 bison-cattle hybrid (*34*), after running ancesty_HMM with the parameters ‘-e 0.02 -a 2 0.5 0.5 -p 0 10000 0.5 -p 1 1 0.5’ (fig. S16). Given the limited sample size of high coverage ancient bison genomes in our bison ancestry source panel, we then filtered this putative set of AIMs by removing sites where any of 99 ancient bison genomes older than 1,000 years had reads carrying the cattle allele, or where fewer than 11 of these ancient bison had any reads. This gave a final set of 5,358,142 AIMs for use in local ancestry inference.

We used SAMtools mpileup (-B -q25 -Q30) to generate generate read piles AIM sites for each individuals, and then modeled local ancestry across the genome for each modern bison using ancestry_HMM with the parameters '-e 0.02 -a 2 0.99 0.01 -p 0 100000 0.99 -p 1 -30 0.01'. This models a 1% cattle ancestry pulse occurring 30 generations in the past, though allows admixture timing to be estimated from the data. Ancestry posterior probabilities were then converted into hard ancestry calls, requiring a minimum posterior probability of 90%. Adjacent ancestry calls were merged into tracts, requiring a minimum of 10,000 AIMs to call a segment. We used a weighted blocked jackknife (*146*) in 1 megabase windows to calculate standard errors for our estimates of the overall proportion of cattle ancestry. We implemented the jackknife using code based on the jackknife.R script (https://github.com/simonhmartin/genomics_general). For bison that were identified with significant evidence of cattle ancestry, we performed an additional local ancestry inference analysis with the same ancestry_HMM model using 100 bootstrap replicates to assess the distribution of inferred admixture dates.

Altogether, we identified cattle ancestry in 32 of the 97 modern bison examined in this study (Fig. 5C). When present, the proportion of cattle introgression was low, generally less than 1% (Fig. 5C). Cattle ancestry was found almost exclusively in large, heterozygous segments (fig.

S17), consistent with a recent inferred date of admixture (Fig. 5E). Given that cattle ancestry is rare and in a heterozygous state, it is possible that it will be lost over time through drift, especially if gene flow between herds is sufficient to prevent the fixation of particular segments of cattle ancestry within individual herds. To highlight this process, we compared the cattle ancestry tracts between two parent-offspring pairs from the FTN herd (fig. S18).

To test for whether cattle ancestry was distributed non-randomly across the genomes of these individuals, we used a permutation approach. In order to avoid overcounting segments that were inherited from a very recent common ancestor, we first pruned pairs of up to second-degree relatives identified using READv2 (as described in 'Pruning relatives') by randomly removing one individual within each pair of relatives until no related individuals remained. We then shuffled all tracts present among individuals in the pruned dataset randomly across the genome, and split the cattle genome into 1 Mb windows and counted for both the empirical and permuted data the number of cattle segments overlapping each window using BEDtools (*147*) intersect (fig. S19A). We repeated this pruning and shuffling process 1,000 times, and, to test for cattle ancestry hotspots, we calculated an empirical $p$-value for each 1 Mb window as the proportion of times the permuted data had a count of overlapping cattle segments as high or higher than the empirical data (fig. S19B). Finally, we also compared the overall distribution of overlap counts across all windows between the empirical and permuted data (fig. S19C), and tested whether these distributions differed using a Chi-squared test (performed with the chisq.test R function with simulate.p.value = TRUE), with marginally significant evidence supporting that they were distinct (Chi-squared $p = 0.016$). We observed that the empirical distribution had more segments without cattle ancestry along with a longer tail, perhaps suggesting the presence of both cattle ancestry deserts and hotspots. However, given the recent and limited nature of the interaction between bison and cattle, and the low sample size of individuals with cattle ancestry, it remains unclear whether these patterns can best be explained by demography, recombination, or selection.

As we only had four high coverage ancient genomes, and these were geographically and temporally removed from modern bison, we wanted to evaluate whether this limited sample size was sufficient for representing unadmixed bison ancestry in modern bison. We constructed a modern source panel consisting of 20 high coverage bison genomes, which were shown to be largely free from cattle admixture using both our local ancestry inference framework and *f4*-statistics of the form *f4(modern bison, Horner_WY_10k; angus, gaur*). The modern bison source panel was constructed using the same filtering procedure as the ancient bison source panel described above (without filtering using the low-coverage ancient bison read-level data) and then local ancestry inference was repeated using the modern source panel on all modern bison which were not members of the panel. Using this larger panel of modern bison instead of the four steppe bison provided highly concordant results (fig. S20), suggesting that even a few high coverage ancient bison are sufficient as an ancestry source panel, likely due to the millions of years of divergence separating bison and cattle (38).

Our estimates of the extent of cattle ancestry in modern bison appear to be substantially smaller, both within and across individuals, than those that have been previously published, most comprehensively in Stroupe *et al. (20)*. We sought to understand why this might be by comparing our local ancestry inference results to analyses in Stroupe *et al.* for the 24 individuals that had publicly available data (fig. S21). All of these bison were inferred by Stroupe *et al.* to have cattle ancestry, although we recover cattle ancestry tracts in only 7 out of 24 of the individuals. While the exact coordinates of inferred cattle ancestry tracts from previous analyses were not available, comparing qualitatively, the majority of segments identified in our analysis were also found in Stroupe *et al.*, such that the regions of cattle ancestry we identified were largely a subset of those previously published. Segments present in Stroupe *et al.* but not in our analyses occurred almost entirely in regions with very few ancestry informative markers (Fig. 5F). Examples of this phenomenon can be found in large tracts with very low AIM density on chromosomes 10 (at ~20 Mb), 12 (at ~70 Mb), and 23 (at ~25 Mb), though it is also evident for shorter tracts. This suggests that these regions do not arise from cattle introgression, but are instead areas of low differentiation between bison and cattle. In some cases, these regions are poorly resolved in the cattle assembly (including the highlighted region on chromosome 12), and so bioinformatic artifacts may inflate similarity between bison and cattle and lead to false positives. Consistent with this, poorer quality samples, including historical samples with shorter read lengths such as HistW_1937/W13, tended to have a higher number of short segments of cattle ancestry in Stroupe *et al*. Such segments were not found in our local ancestry analysis. As one of the methods used in Stroupe *et al.* does not distinguish between admixture and other explanations for genomic similarity (*148*), like incomplete lineage sorting, we infer that previous analyses may be complicated by disentangling admixture from other processes. Using data from ancient bison therefore allows for detecting admixture with greater sensitivity.

**Supplementary Figures**

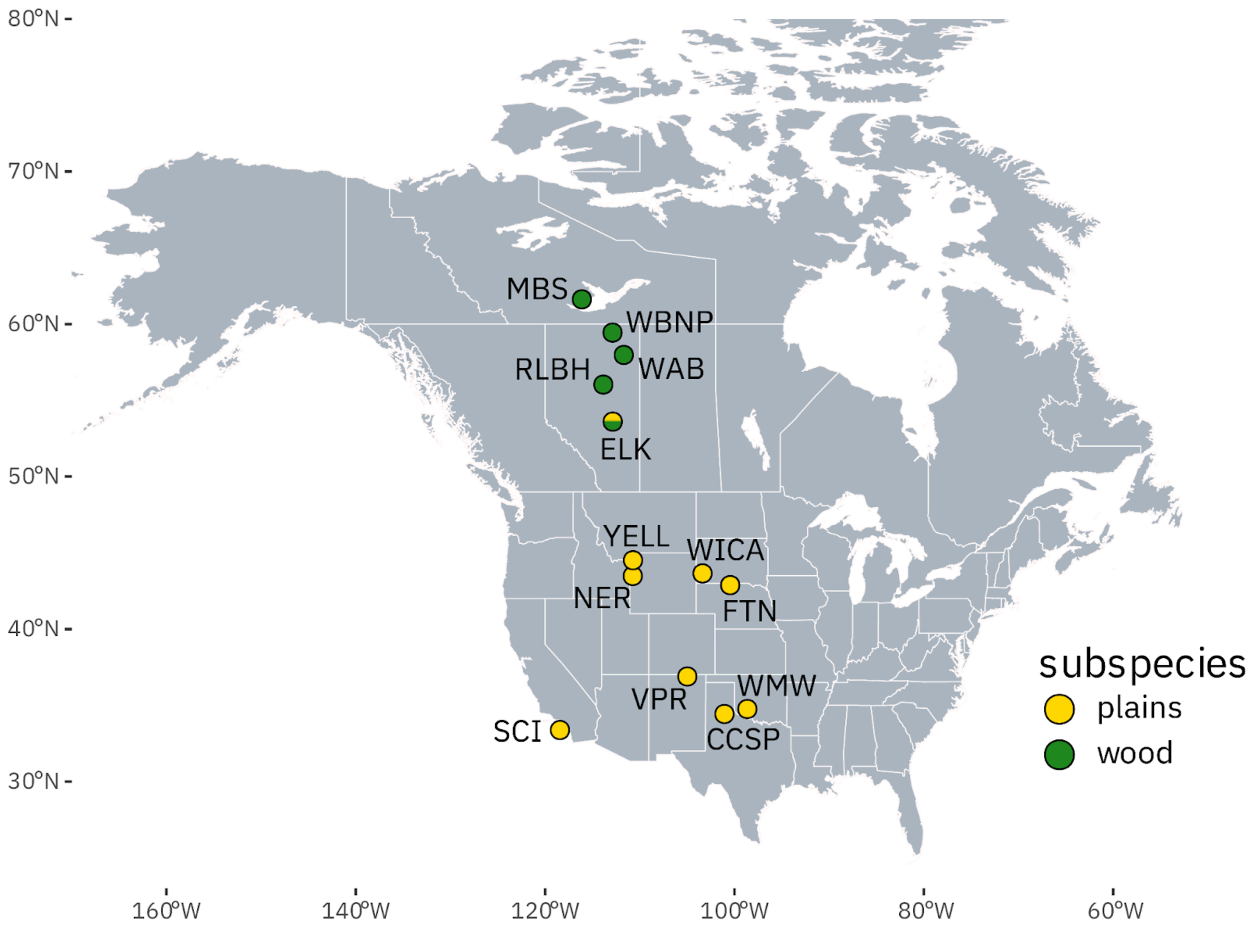


**Figure S1. Locations of key modern bison herds used in this study.** Wood bison herds are shown in green, while plains bison are in yellow. Elk Island contains both plains and wood bison which are kept geographically isolated within the park. Herd abbreviations: CCSP = Caprock Canyons State Park; SCI = Santa Catalina Island; NER = National Elk Refuge/Grand Teton National Park; FTN = Fort Niobrara National Wildlife Refuge; VPR = Vermejo Park Ranch; WICA = Wind Cave National Park; WMW = Wichita Mountain Wildlife Refuge; YELL = Yellowstone National Park; ELK = Elk Island National Park; MBS = Mackenzie Bison Sanctuary; RLBH = Ronald Lake Bison Herd; WAB = Wabasca; WBNP = Wood Buffalo National Park.

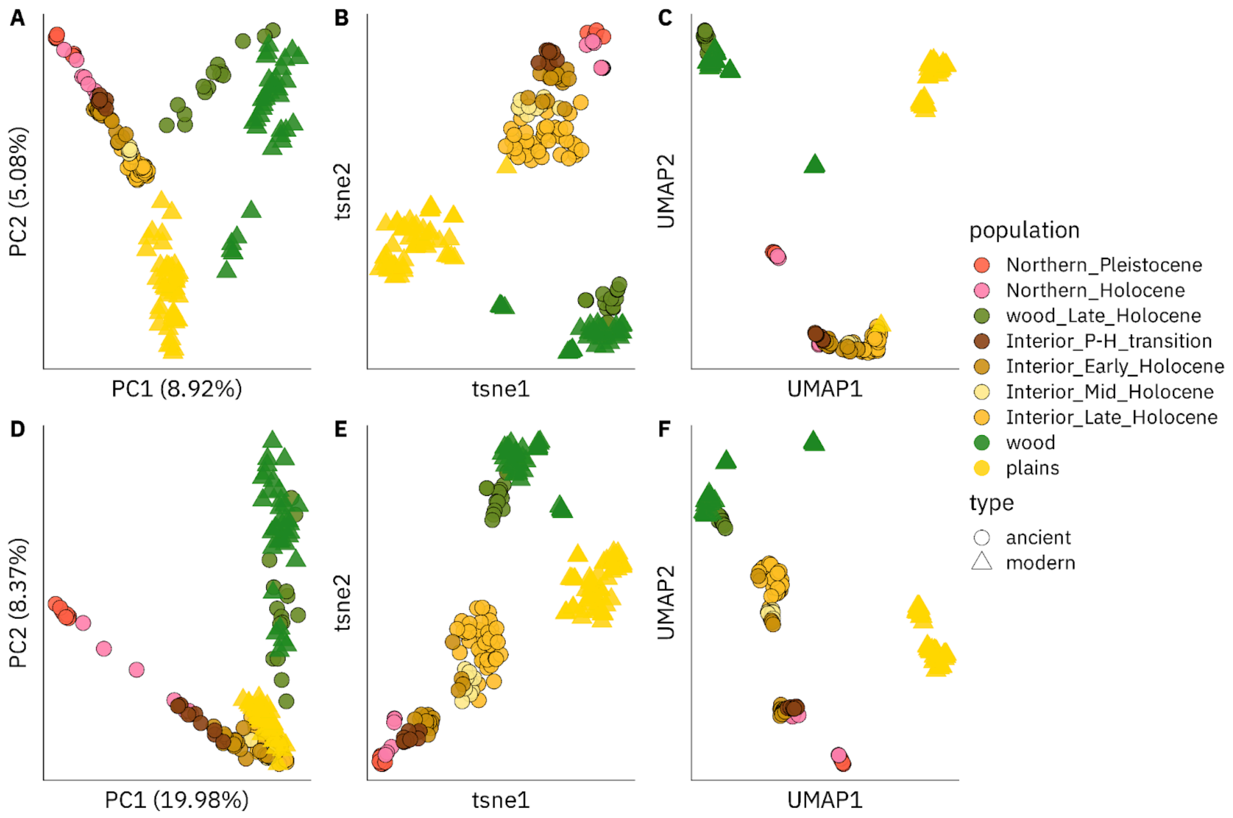


**Figure S2. f-statistic based dimensionality reduction analyses for post-Last Glacial Maximum North American bison.** For all bison examined in this study, *f4*- (A-C) and *f3*-statistic (D-F) based dimensionality reduction analyses: PCA (A,D), t-SNE (B,E), UMAP (C,F). Matrices of all possible combinations of *f4- or f3*-statistics relating each individual in the dataset were used as input. Bison are colored by temporal, geographic, and genetic (for ancient wood bison) groupings, with ancient samples shown in circles and modern samples using triangles.

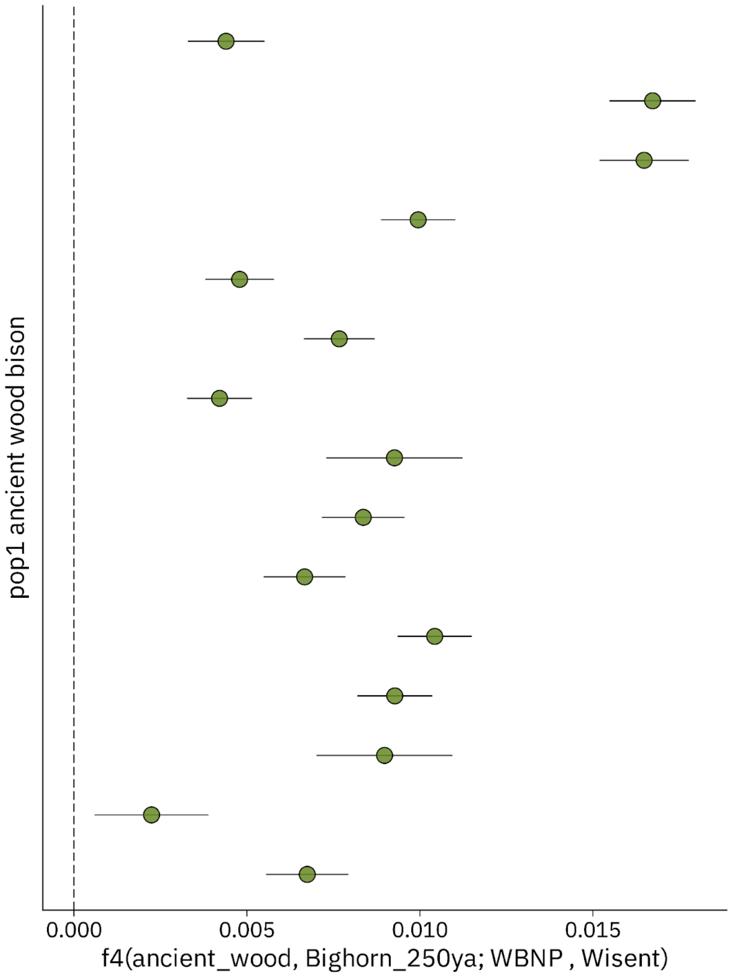


**Figure S3. Allele sharing between ancient and modern wood bison.** *f4*-statistics testing for affinity between ancient and modern wood bison, relative to late Holocene bison from Wyoming (Bighorn_250ya). Each point represents a different ancient wood bison. Error bars represent ±3 standard errors.

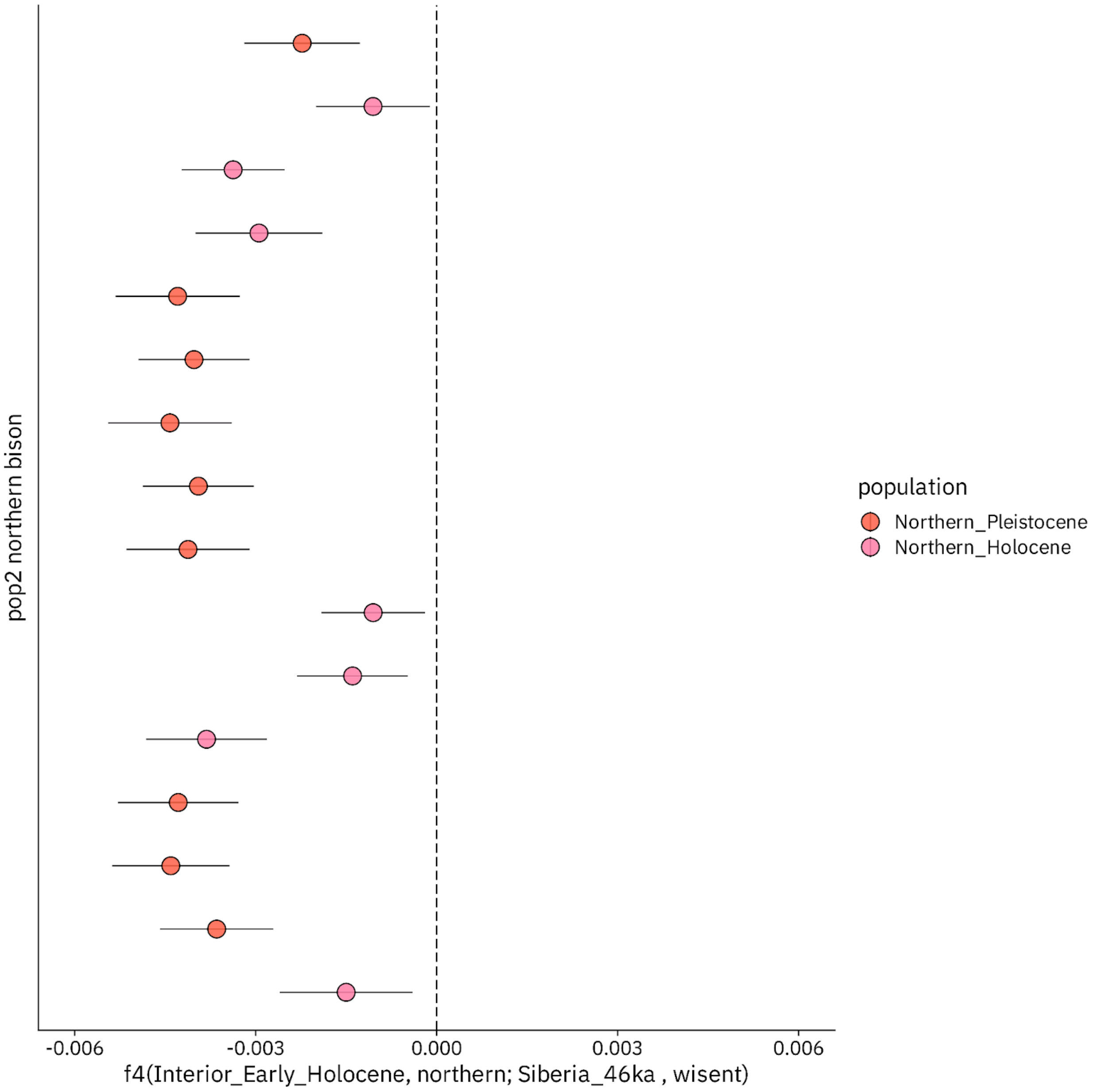


**Figure S4. Siberian-related ancestry in northern bison.** *f4*-statistics testing for affinity between remnant, post-Last Glacial Maximum northern bison and a Late Pleistocene Siberian steppe bison (Siberia_46ka), relative to Interior Early Holocene bison (~10ka). All of these northern bison have significant allele sharing with Siberian bison, suggesting Siberian-related ancestry not found in Interior bison is retained in these populations after the LGM. Each point depicts a different ancient northern bison used in the comparison. Error bars represent ±3 standard errors.

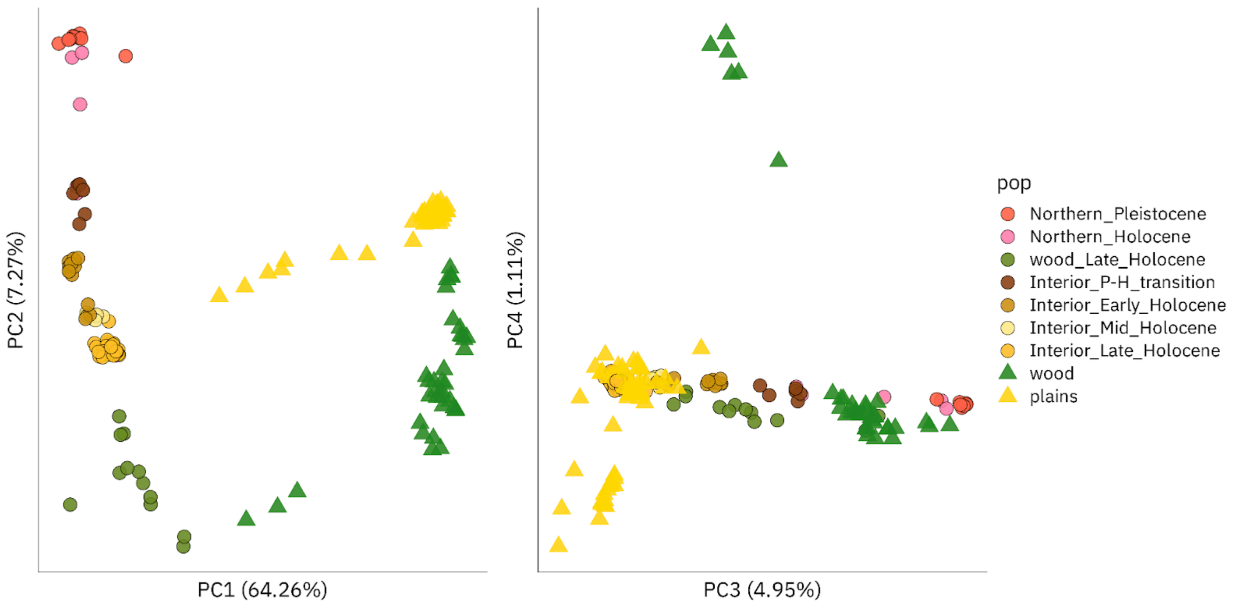


**Figure S5. Genotype likelihood-based principal component analysis of post-Last Glacial Maximum North American bison.** PCA was performed with PCAngsd using genotype likelihoods for transversion variants across all 172 bison with ≥0.5-fold average coverage. Population labels are the same as those in Fig. S2.

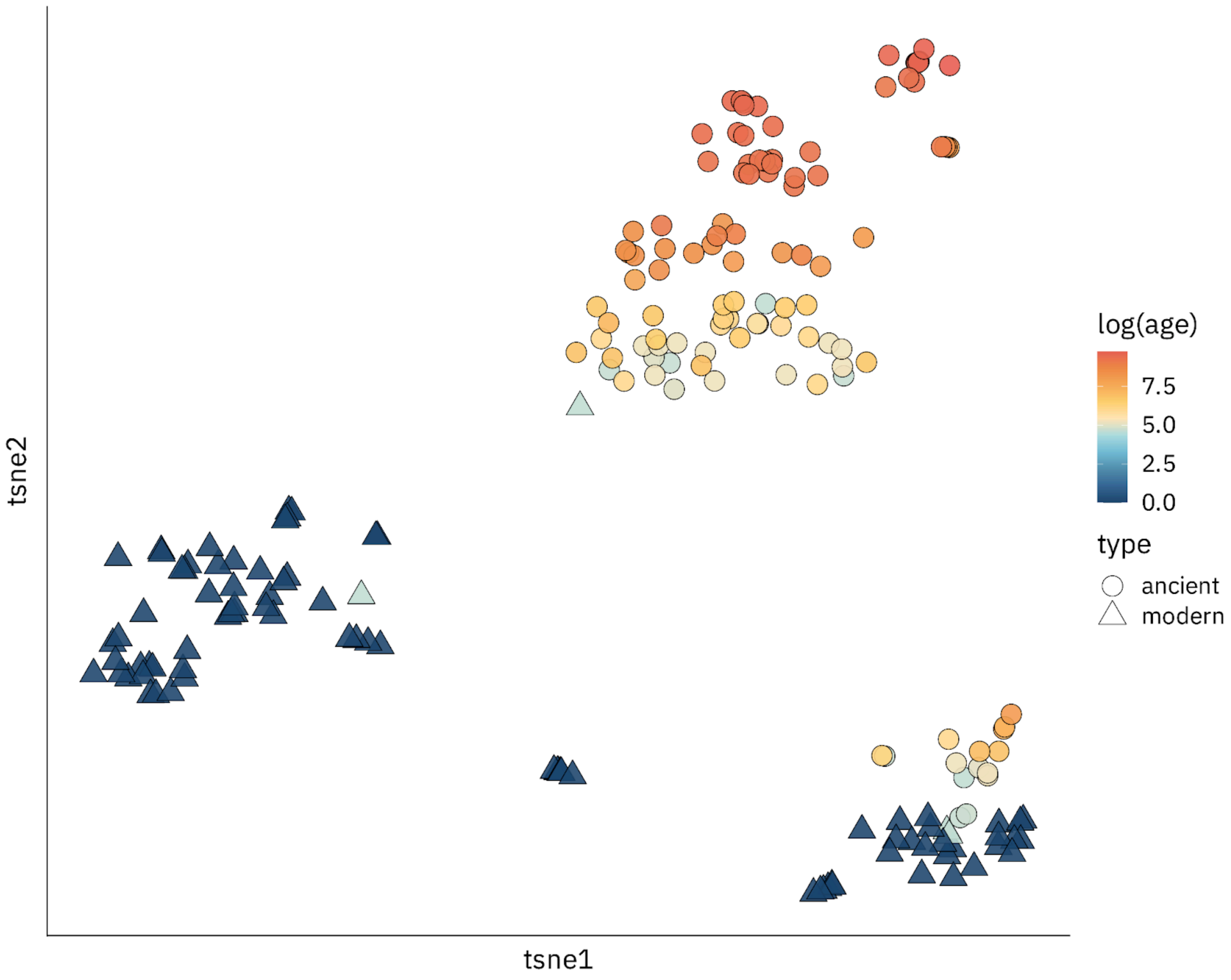


**Figure S6. Genetic relationships by time.** *f4*-based t-SNE analysis for all post-LGM bison, colored by log(sample age). The ages of all modern individuals were set to 0.

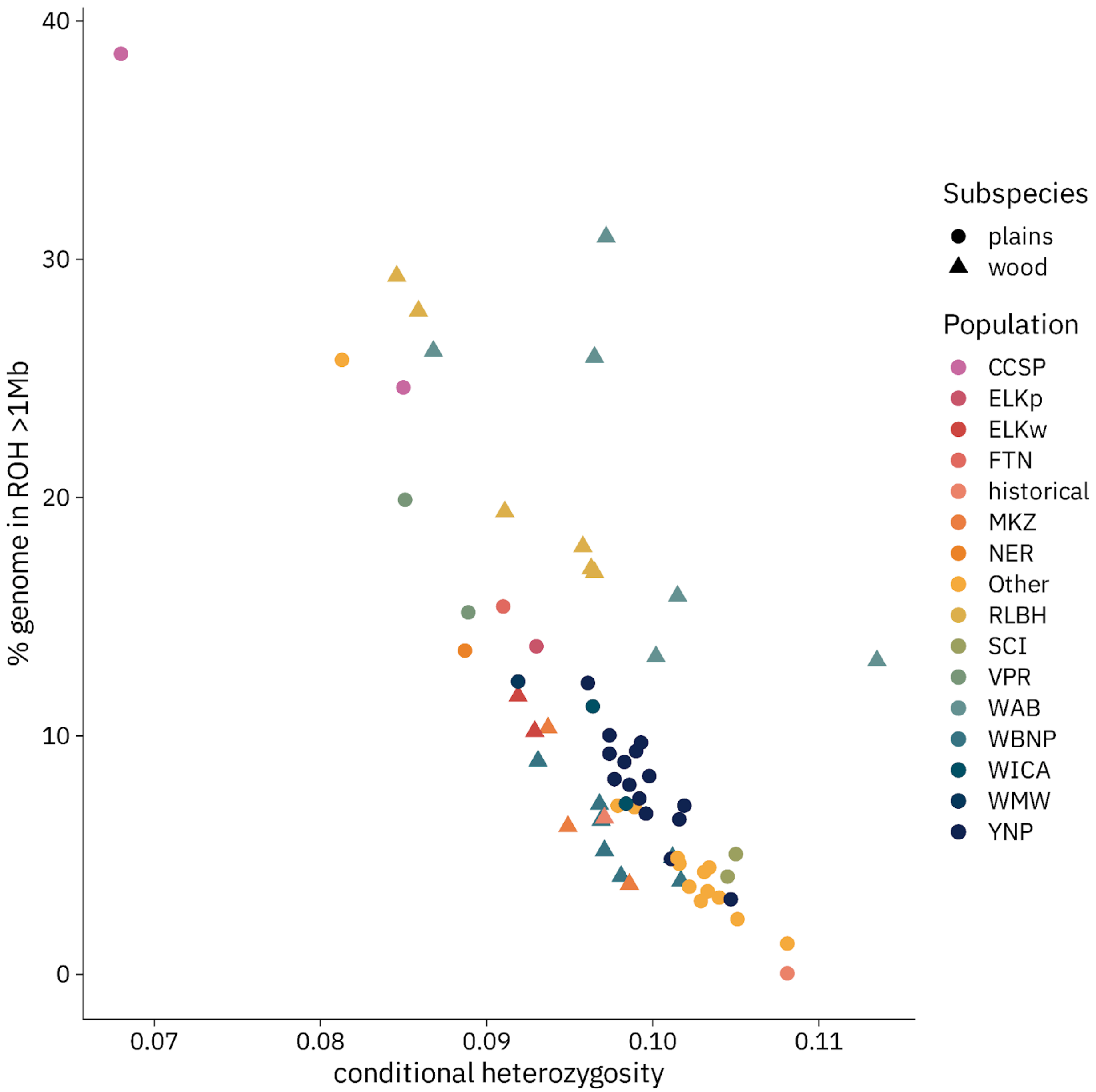


**Figure S7. Relationship between genetic diversity and inbreeding in modern bison.** Genetic diversity was estimated as pseudodiploid conditional heterozygosity estimates by randomly sampling two reads. The proportion of the genome contained in ROH was estimated using ROHan for 1 Mb windows.

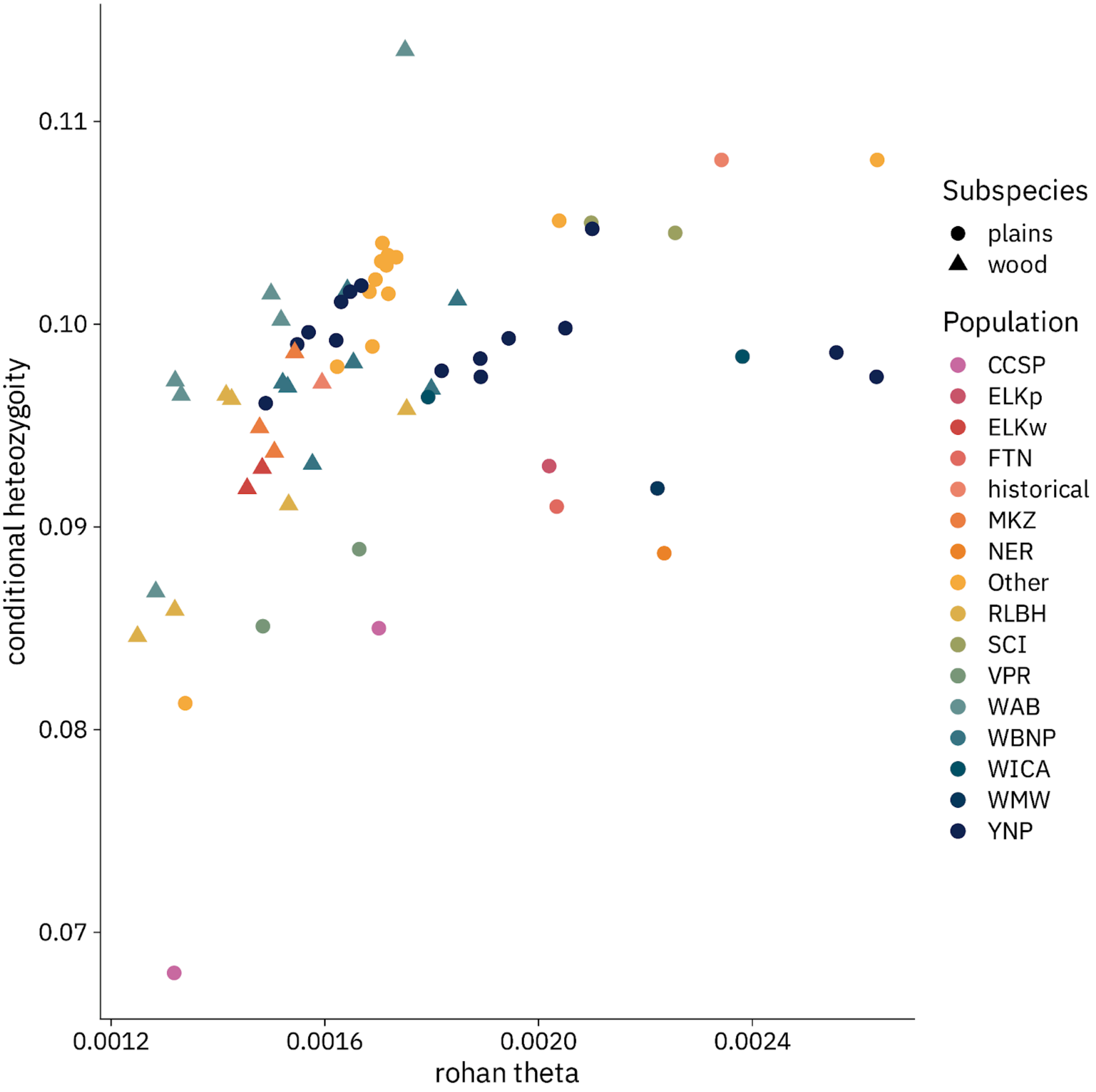


**Figure S8. Comparing different estimates of heterozygosity in modern bison.** Whole-genome estimated theta (including runs of homozygosity) calculated from ROHan compared to conditional heterozygosity estimated through pseudodiploid sampling at 3.6M transversions ascertained from heterozygous sites in 7 high coverage bison genomes.

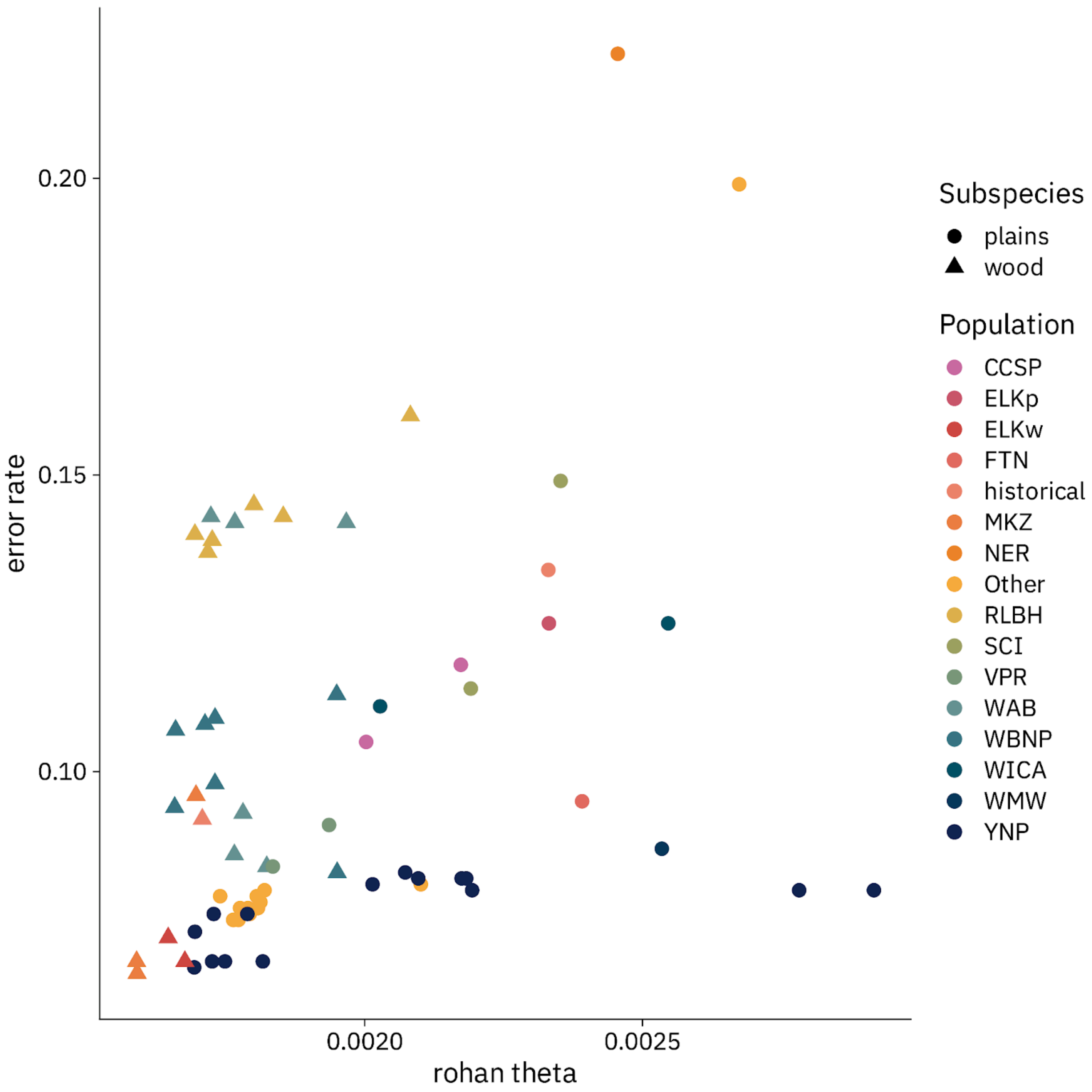


**Figure S9. Comparing ROHan heterozygosity estimates and per-sample error rates in modern bison.** Theta was estimated using ROHan excluding regions in ROH. As a proxy for per-sample error rates, we used the rate of non-biallelic bases appearing at 3.6M transversions ascertained from heterozygous sites in 7 high coverage bison genomes. Only individuals with ≥10,000 SNPs covered by at least two reads are shown.

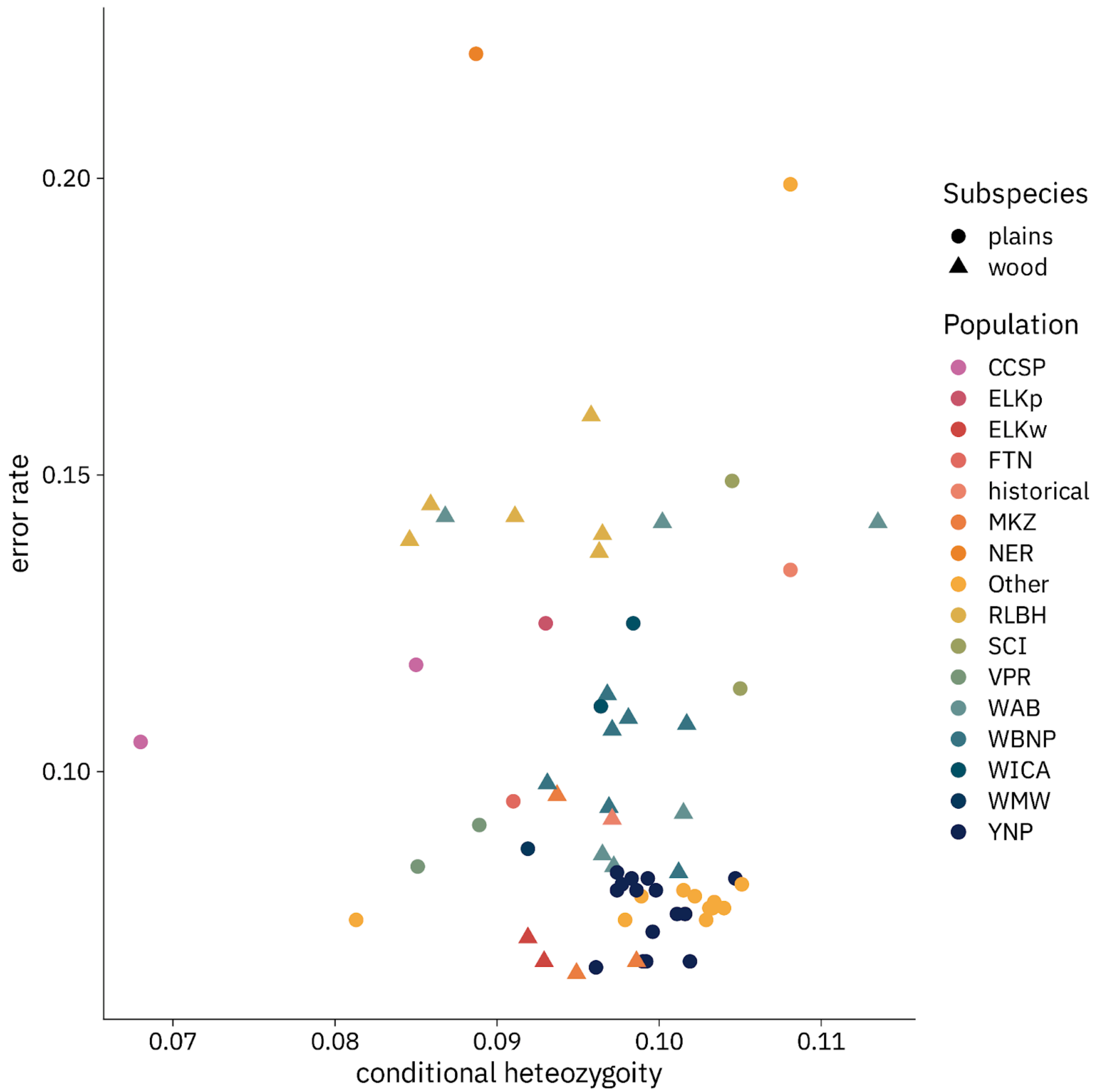


**Figure S10. Comparing conditional heterozygosity and per-sample error rates in modern bison.** Conditional heterozygosity was estimated through pseudodiploid sampling of reads at 3.6M transversions ascertained from heterozygous sites in 7 high coverage bison genomes. As a proxy for per-sample error rates, we used the rate of non-biallelic bases appearing at the same set of transversions. Only individuals with ≥10,000 SNPs covered by at least two reads are shown.

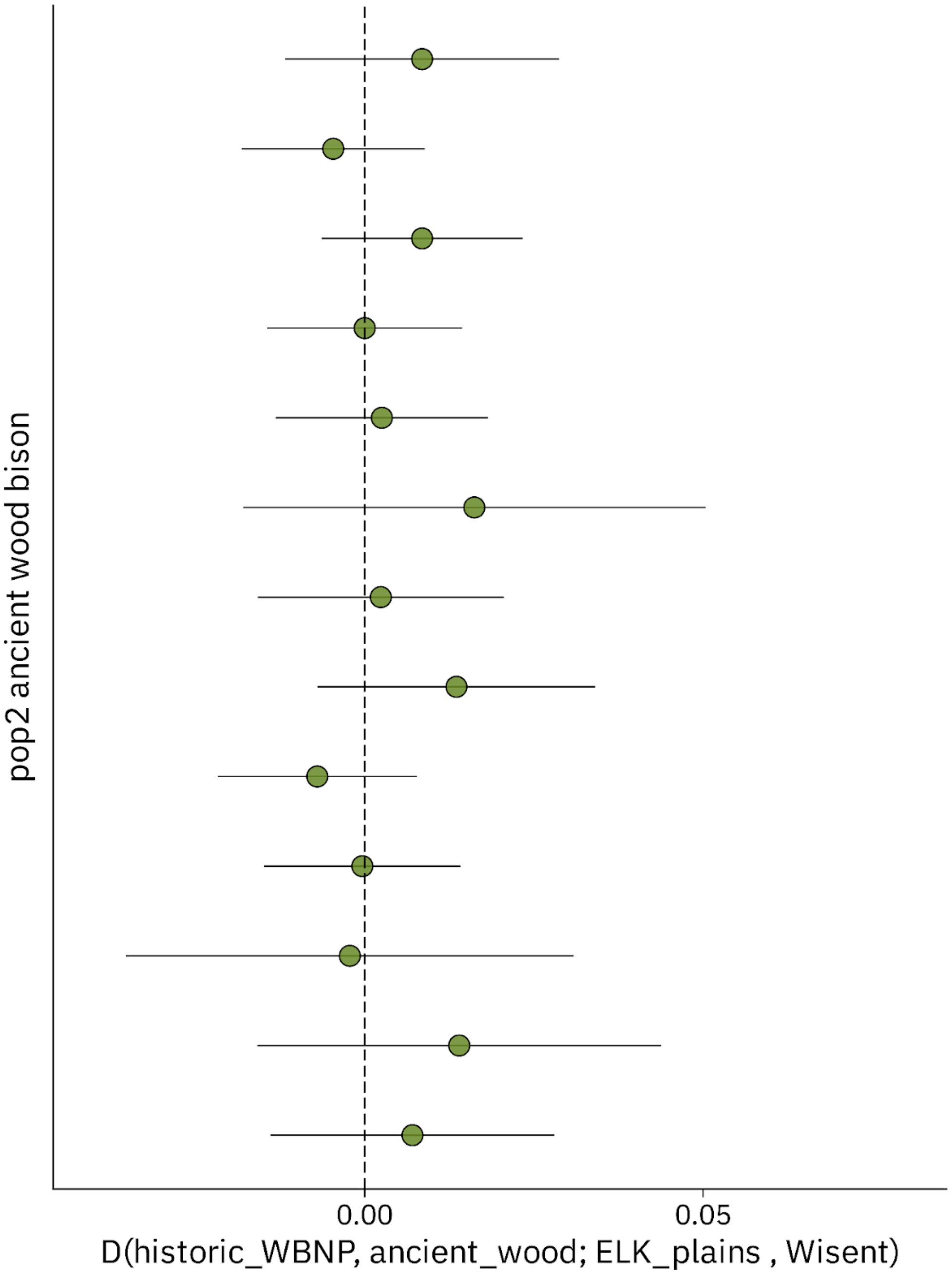


**Figure S11. Allele sharing between ancient wood bison individuals and Elk Island plains bison, relative to historical WBNP.** *D*-statistics testing for affinity between ancient wood bison and modern Elk Island plains bison, relative to historical WBNP bison. Each point depicts a different ancient wood bison. Error bars represent ±3 standard errors.

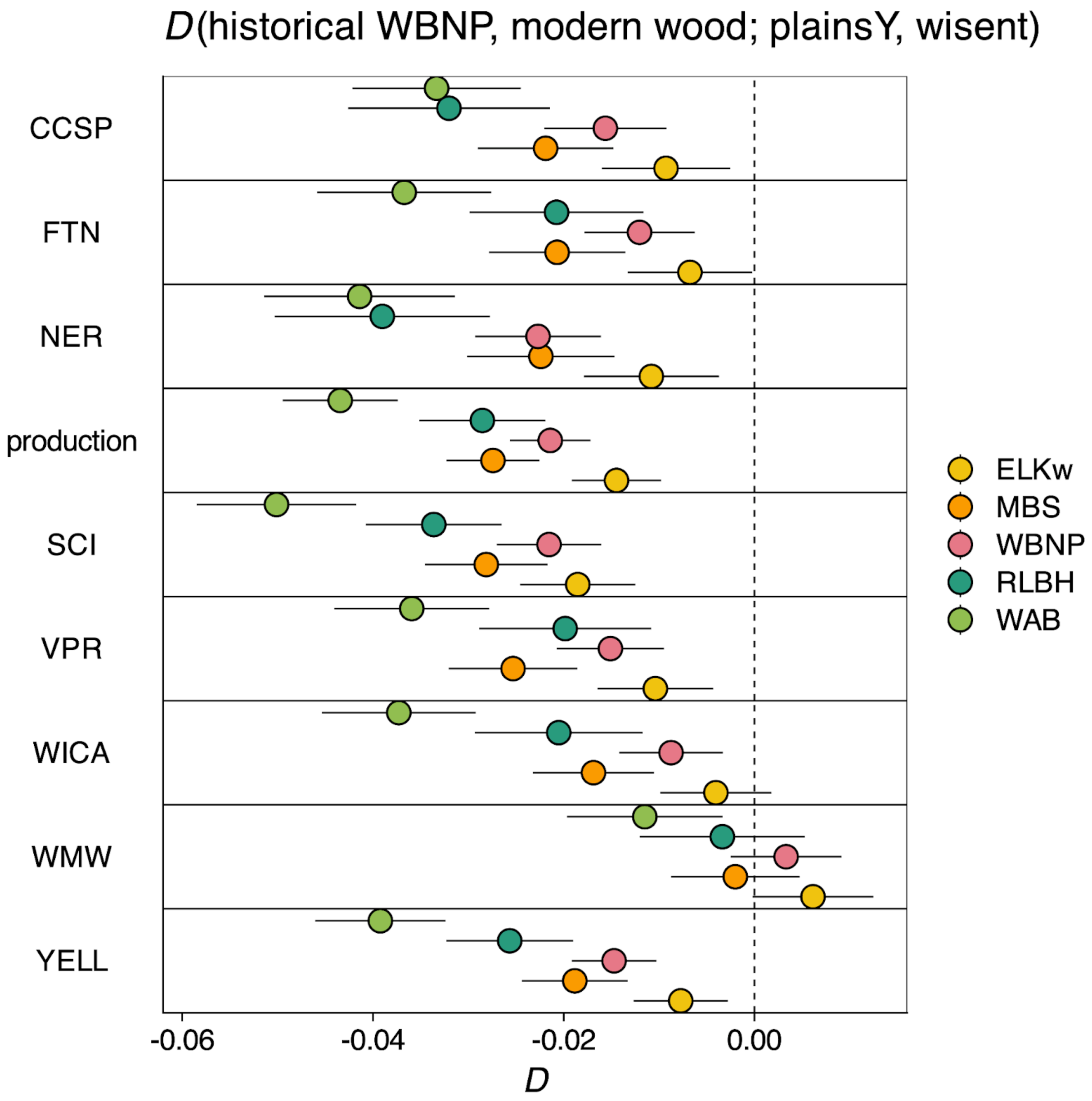


**Figure S12. Evidence of plains bison ancestry in wood bison regardless of source plains herd.** *D*-statistics testing for differential allele sharing between the five modern wood bison herds (colors) and different non-Elk Island plains bison sources (y-axis), relative to two ~175 year old WBNP bison. All modern wood bison share more alleles with plains bison relative to these historical WBNP bison, regardless of the plains bison herd used in the comparison. Error bars represent ±3 standard errors.

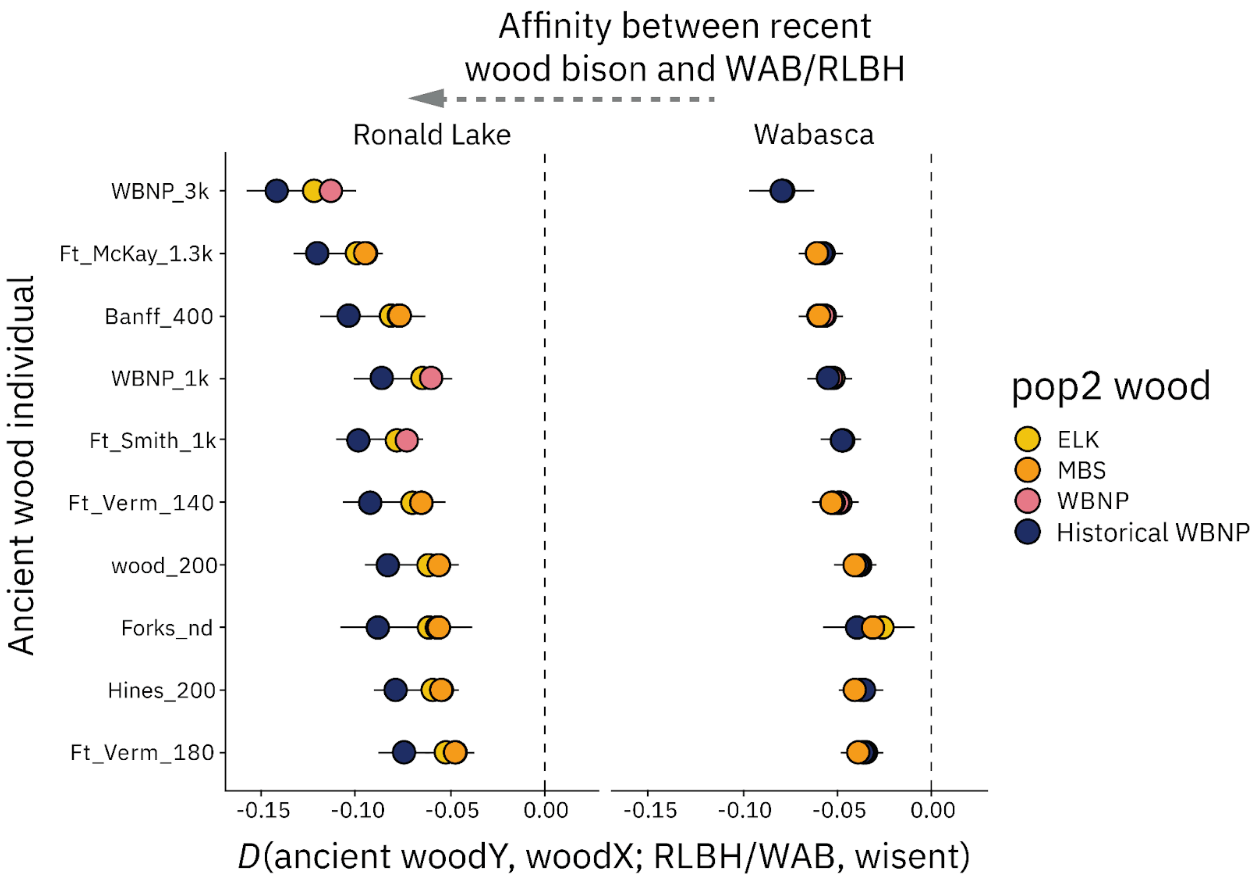


**Figure S13. Wabasca and Ronald Lake share a close relationship with other modern wood bison herds.** Testing for differential allele sharing between the Ronald Lake (left) and Wabasca (right) wood bison herds to either ancient wood bison or modern wood bison herds. Different ancient wood bison used in the comparison are shown on the y-axis, with each point representing a different modern or historical wood bison herd. Both Ronald Lake and Wabasca share more alleles with recent wood bison than with any ancient wood bison individual. Error bars represent ±3 standard errors.

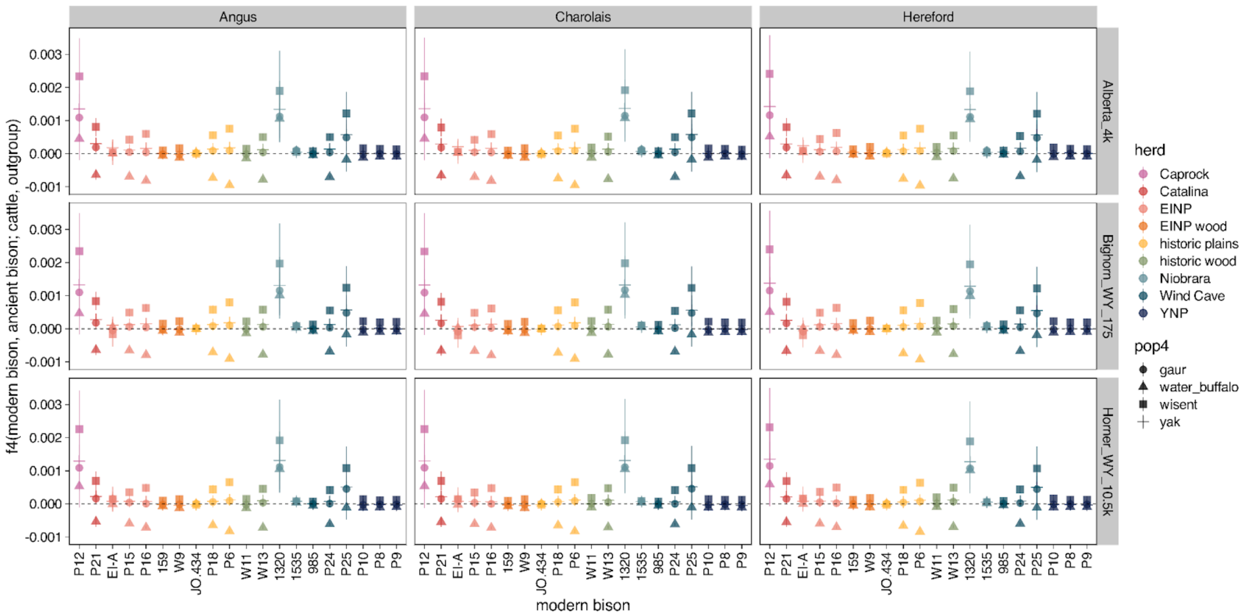


**Figure S14. Testing for cattle introgression in modern bison using alignments to the bison genome and bovine-ascertained variants.** *f4*-statistics testing for allele sharing with cattle in 20 modern bison from 9 different herds, using different cattle breeds (columns), ancient bison (rows), and outgroups (symbols). Statistics were calculated with pseudohaploid genotypes called for variants ascertained in a range of bovine species: bison, cattle, gaur, yak, wisent, and water buffalo. Error bars represent ±3 standard errors.

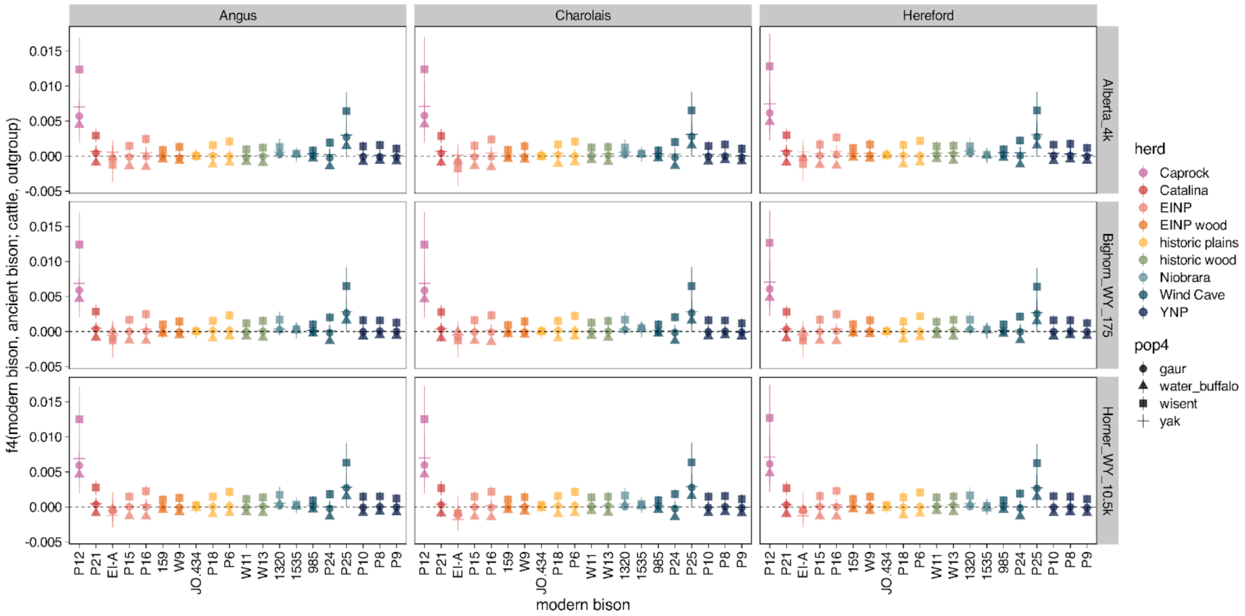


**Figure S15. Testing for cattle introgression in modern bison using alignments to the bison genome and bison-ascertained variants.** *f4*-statistics testing for allele sharing with cattle in 20 modern bison from 9 different herds, using different cattle breeds (columns), ancient bison (rows), and outgroups (symbols). Statistics were calculated with pseudohaploid genotypes called for variants ascertained in 10 modern bison. Error bars represent ±3 standard errors.

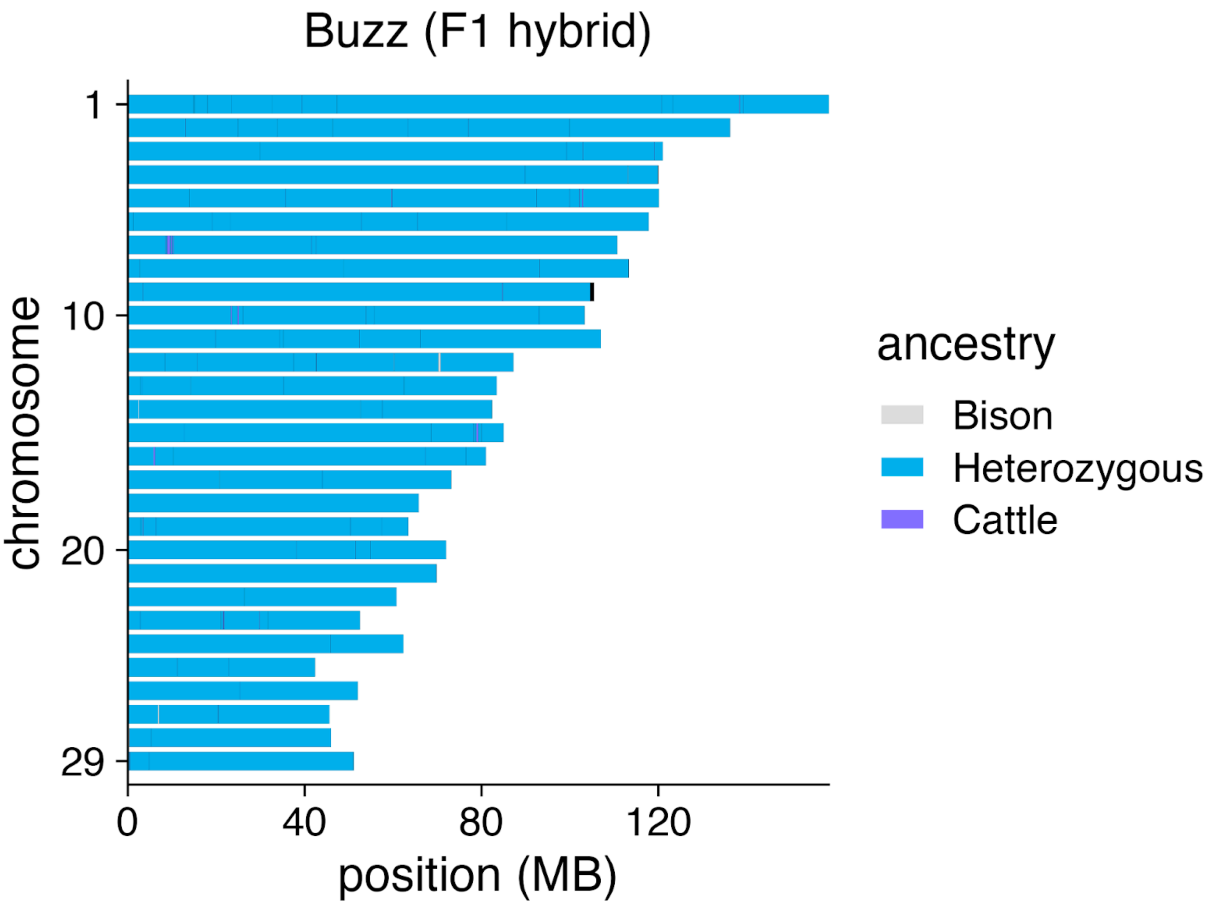


**Figure S16. Local Ancestry Inference for a bison-cattle F1 hybrid.** Inferred diploid ancestry state for 'Buzz', a F1 Yellowstone bison-Simmental cattle cross (*34*). Ancestry state is shown for all 29 autosomes. 99.87% of Buzz's genome was correctly inferred to be heterozygous for bison and cattle ancestry. Results are shown prior to filtering, and markers which were not confidently assigned as heterozygous were removed for subsequent LAI analyses.

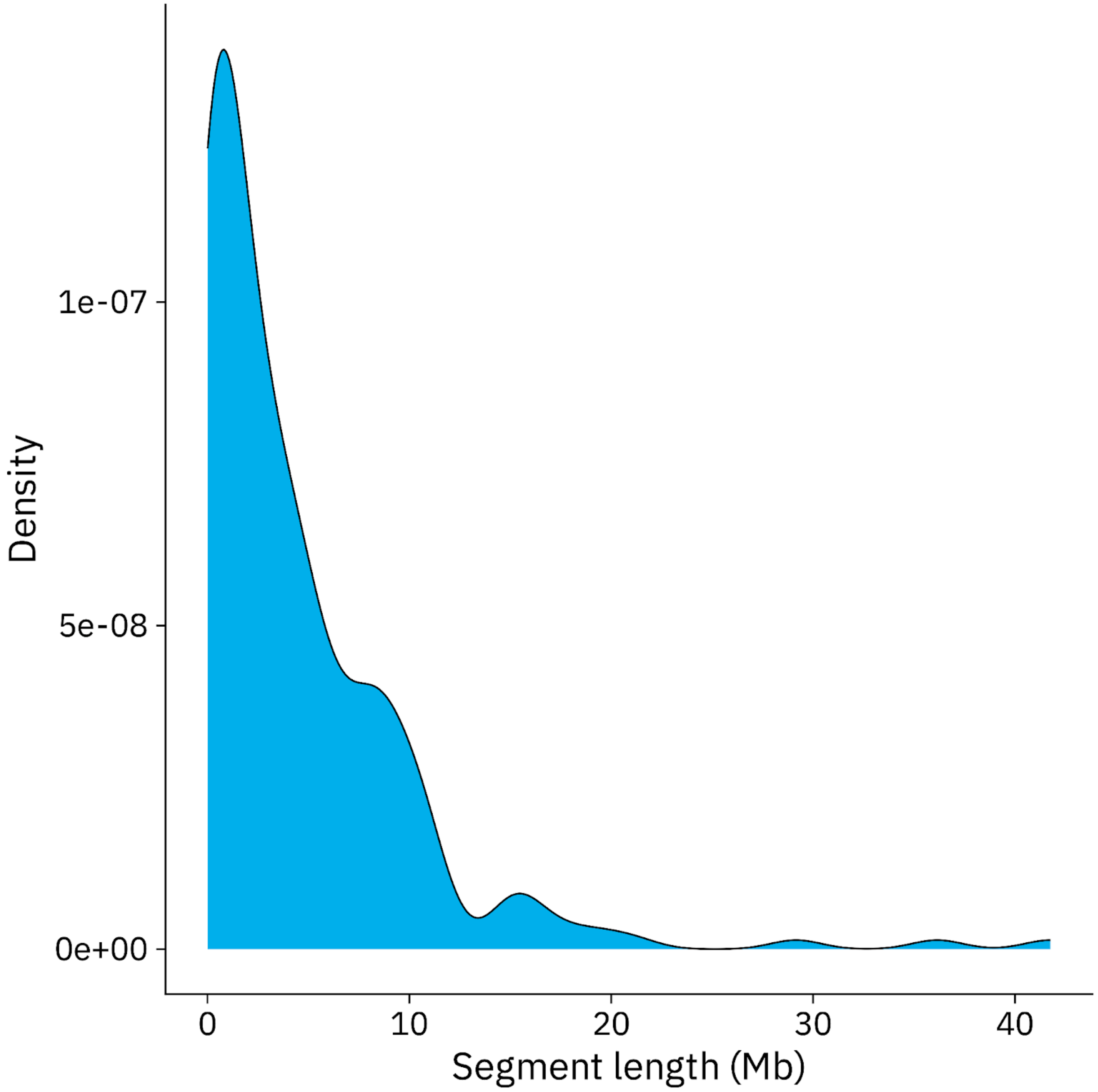


**Figure S17. Density of cattle ancestry tract lengths.** Tracts are combined across all sequenced modern bison, identified using local ancestry inference performed with steppe bison and modern cattle ancestry source panels.

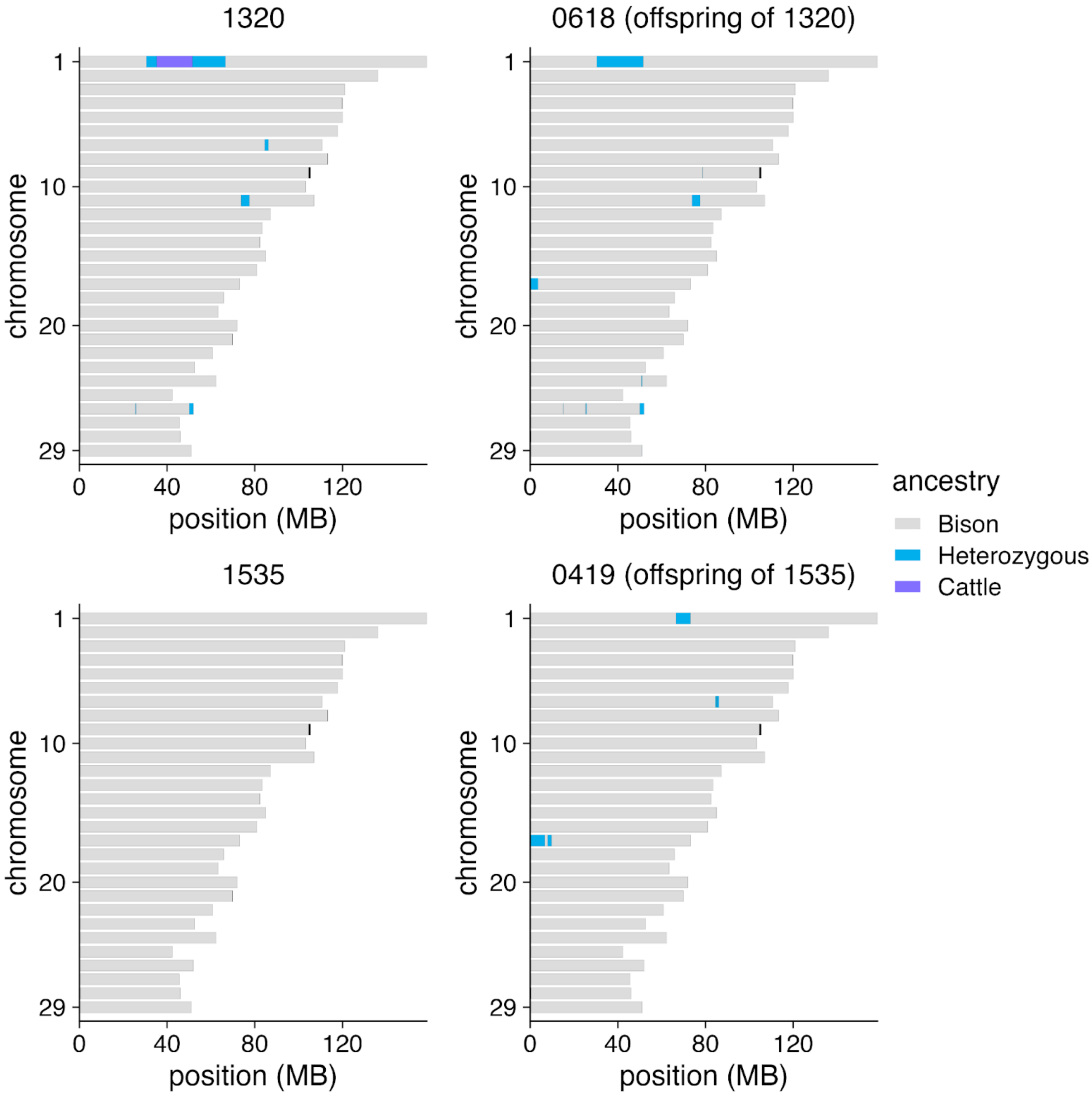


**Figure S18. Cattle ancestry comparisons for two parent-offspring pairs.** Inferred diploid autosomal ancestry for two parent-offspring pairs from the FTN herd. Local ancestry inference was performed with steppe bison and modern cattle ancestry source panels.

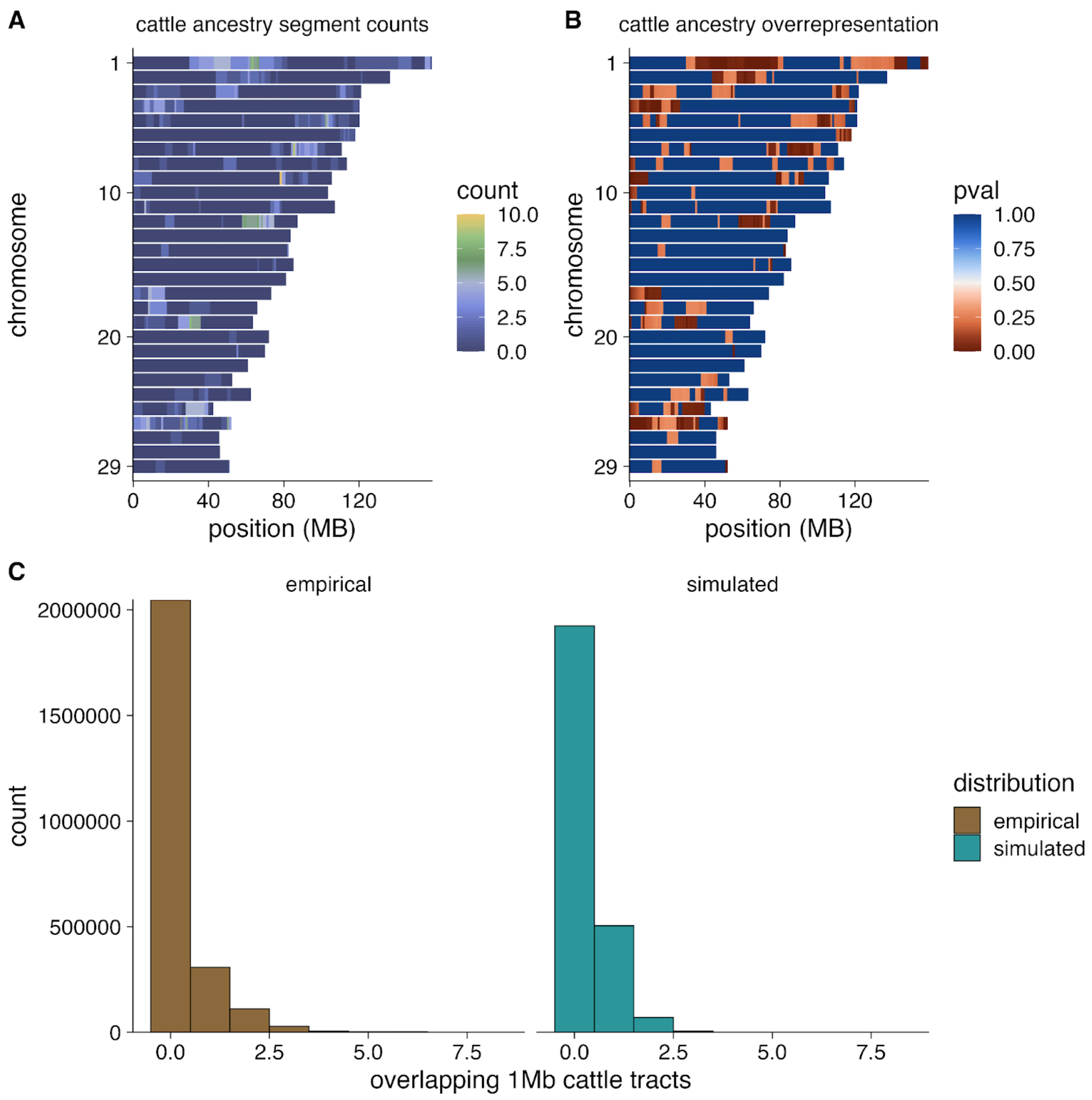


**Figure S19. Distribution of cattle ancestry across modern bison genomes. A)** Counts of the number of cattle segments present across all individual bison in 1 Mb windows. **B)** Empirical *p*-values in 1 Mb windows comparing actual counts of overlapping cattle segments across all individuals to counts obtained by randomly permuting segment locations, following the pruning of close relatives. **C)** Comparison of the distributions of empirical and permuted cattle ancestry overlap counts across all 1 Mb windows.

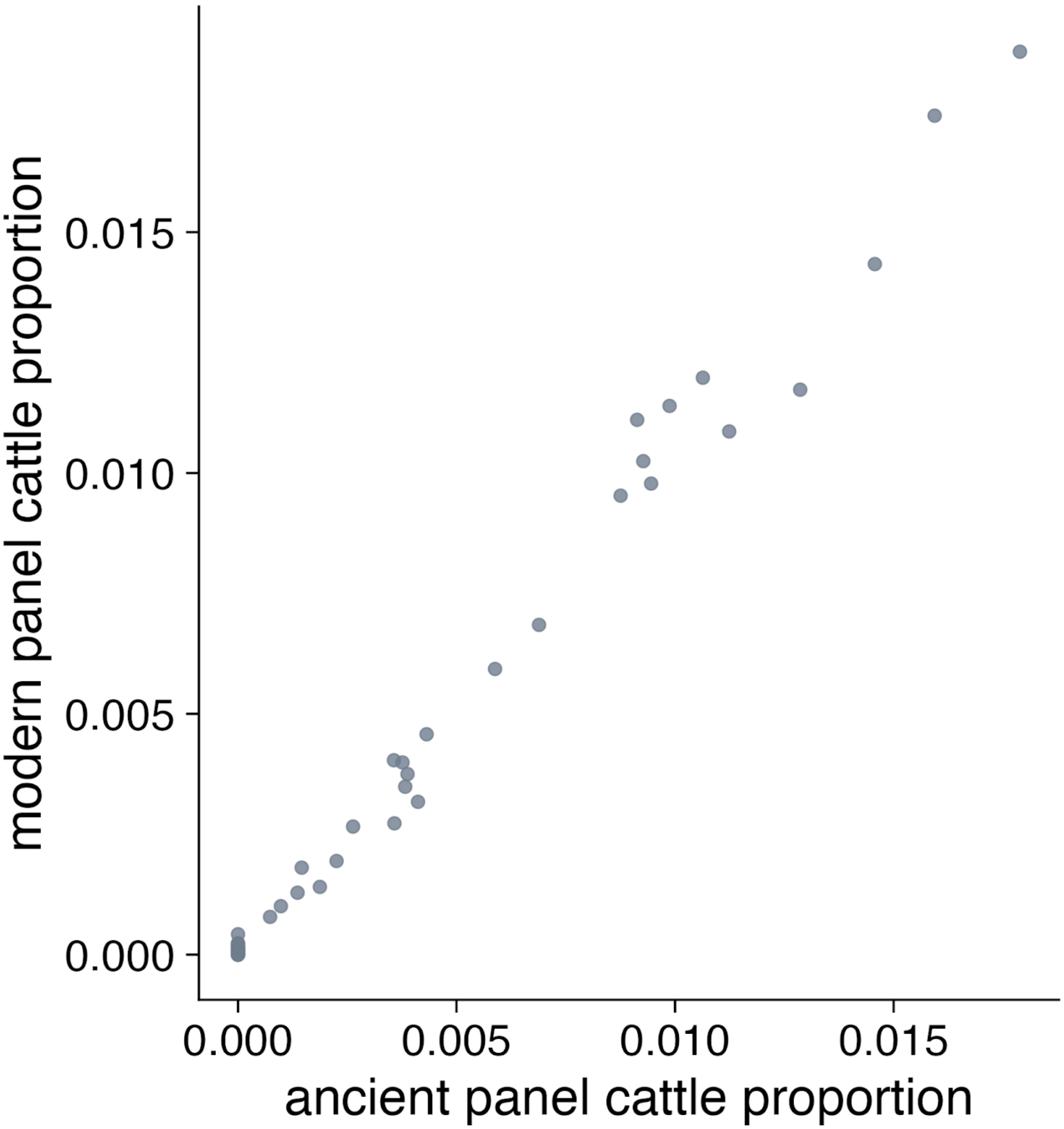


**Figure S20. Inference of cattle ancestry with different bison source panels.** Comparison of overall estimated cattle ancestry proportion using either steppe or modern bison as sources for bison ancestry. Total inferred cattle ancestry proportions are highly similar using either the ancient steppe bison panel or the modern bison panel.

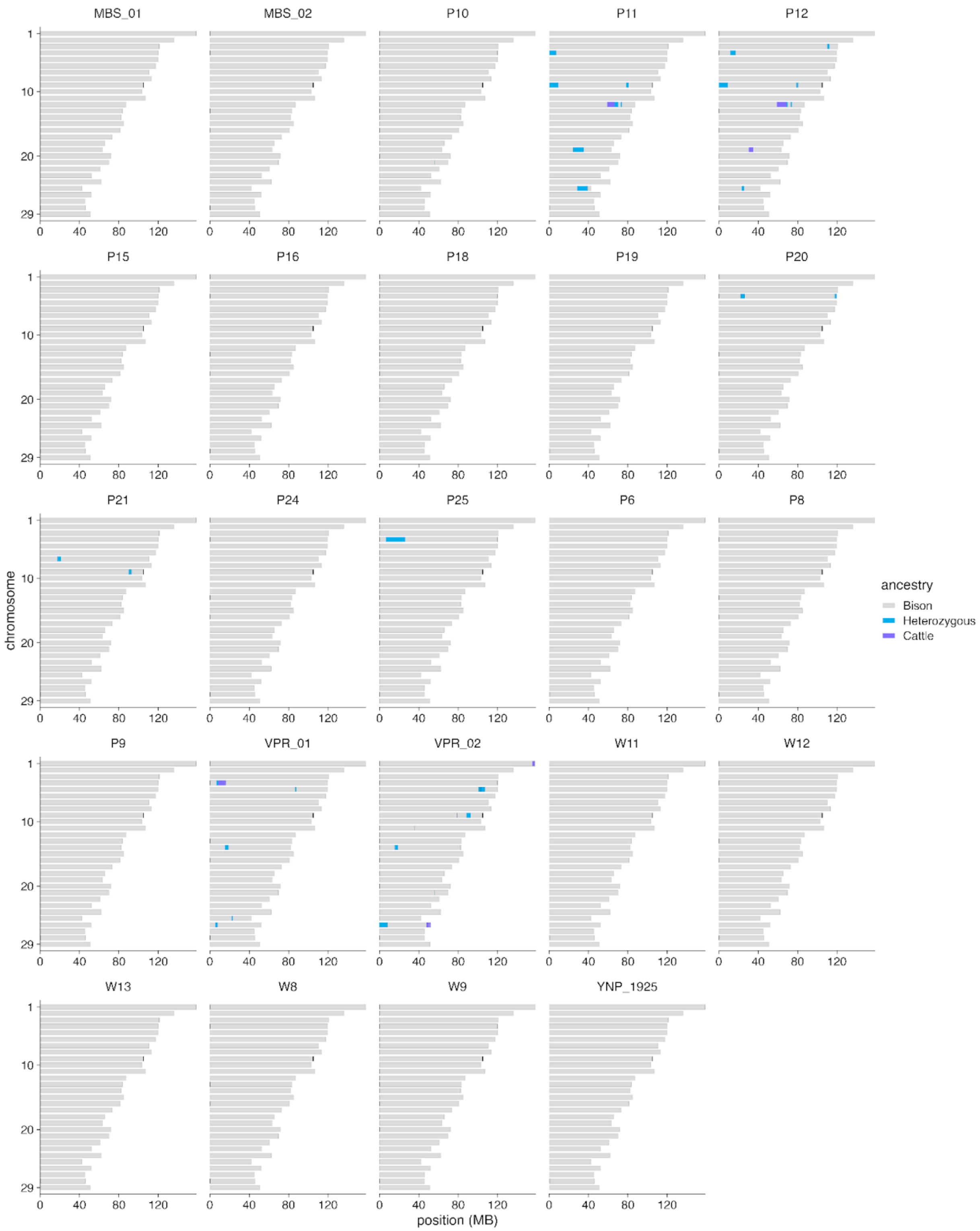


**Figure S21. Inferred local ancestry for previously analyzed individual bison.** Local ancestry inference results for detecting cattle admixture for all individuals published in Stroupe et al. (*20*)**.** Local ancestry inference was performed using ancestry_HMM with steppe bison and modern cattle ancestry source panels.

**Supplementary Tables**

**Table S1.**
**List of samples and metadata for genomes generated in this study** (.xlsx)

**Table S2.**
**List of published bison genomes used** (.xlsx)

**Table S3.**
**Radiocarbon and isotopic data for newly generated radiocarbon dates** (.xlsx)

**Table S4. Pairs of first-degree relatives detected among modern bison.**

| pair | herd |
|---|---|
| 1320,0618 | FTN |
| 1535,0419 | FTN |
| 5164,5180 | NER |
| W3,W2 | ELKw |
| P12,P1 | CCSP |
| P2,P1 | CCSP |
| P2,P12 | CCSP |
| P3,P1 | CCSP |
| P3,P12 | CCSP |
| P3,P2 | CCSP |
| P3,P4 | CCSP |
| P4,P1 | CCSP |
| P4,P2 | CCSP |

**Table S5. Minimum number of admixture events connecting modern wood and plains herds.** Results of qpWave modeling of the relationship between recent wood and plains bison herds.

| f4rank | dof | chisq | p |
|---|---|---|---|
| 4 | 0 | 0.0129 | NA |
| 3 | 2 | 0.542 | 0.76 |
| 2 | 6 | 6.32 | 0.39 |
| 1 | 12 | 14.4 | 0.28 |
| 0 | 20 | 176 | 6.12E-27 |

**Table S6. Modeling modern wood bison ancestry using ancient and modern sources.** Results of qpAdm modeling of the ancestry of modern plains bison herds ("target"), using a rotating approach with the source/reference populations shown in the last 7 columns. All passing models (*p*-value > 0.01 and admixture proportions between 0 and 1) are shown for each target population.

| target | number_of_sources | chisq | p | historical_WBNP | wood_Late_Holocene | ELKp | YELL | WMW | WICA | FTN |
|---|---|---|---|---|---|---|---|---|---|---|
| WBNP | 2 | 4.8635 | 0.1821 | 0.634 | - | 0.366 | - | - | - | - |
| WBNP | 3 | 0.9282 | 0.3353 | 0.631 | - | 0.322 | - | - | 0.047 | - |
| ELKw | 2 | 0.6022 | 0.8959 | 0.745 | - | 0.255 | - | - | - | - |
| ELKw | 2 | 10.2040 | 0.0169 | 0.739 | - | - | 0.261 | - | - | - |
| ELKw | 3 | 0.5031 | 0.4781 | 0.746 | - | 0.231 | 0.023 | - | - | - |
| ELKw | 3 | 0.0390 | 0.8434 | 0.743 | - | 0.244 | - | - | 0.013 | - |
| ELKw | 3 | 0.5646 | 0.4524 | 0.744 | - | 0.244 | - | - | - | 0.011 |
| MBS | 2 | 1.9575 | 0.5813 | 0.641 | - | 0.359 | - | - | - | - |
| MBS | 2 | 11.1861 | 0.0108 | 0.627 | - | - | 0.373 | - | - | - |
| MBS | 3 | 0.3850 | 0.5349 | 0.633 | - | 0.323 | - | 0.044 | - | - |
| MBS | 3 | 0.7743 | 0.3789 | 0.640 | - | 0.302 | - | - | 0.058 | - |
| MBS | 3 | 0.0648 | 0.7991 | 0.641 | - | 0.297 | - | - | - | 0.062 |
| RLBH | 2 | 7.5595 | 0.0560 | 0.454 | - | 0.546 | - | - | - | - |
| RLBH | 3 | 2.0668 | 0.1505 | 0.447 | - | 0.397 | - | - | 0.156 | - |
| RLBH | 3 | 0.5794 | 0.4466 | 0.446 | - | 0.412 | - | - | - | 0.141 |
| WAB | 2 | 10.2741 | 0.0164 | 0.208 | - | 0.792 | - | - | - | - |
| WAB | 2 | 1.3039 | 0.7282 | - | 0.423 | 0.577 | - | - | - | - |
| WAB | 3 | 0.9471 | 0.3305 | 0.206 | - | 0.703 | - | 0.091 | - | - |
| WAB | 3 | 0.3089 | 0.5784 | 0.218 | - | 0.524 | - | - | 0.257 | - |
| WAB | 3 | 0.4316 | 0.5112 | 0.218 | - | 0.531 | - | - | - | 0.251 |
| WAB | 3 | 0.6155 | 0.4327 | - | 0.418 | 0.308 | 0.274 | - | - | - |
| WAB | 3 | 0.7009 | 0.4025 | - | 0.412 | 0.531 | - | 0.057 | - | - |
| WAB | 3 | 0.0637 | 0.8007 | - | 0.423 | 0.542 | - | - | 0.035 | - |
| WAB | 3 | 1.0879 | 0.2969 | - | 0.421 | 0.546 | - | - | - | 0.032 |

**Table S7. Modeling modern wood bison ancestry using only ancient sources.** Results of qpAdm modeling of the ancestry of modern plains bison herds ("target"), using a rotating approach with the source/reference populations shown in the last 5 columns. All passing models (*p*-value > 0.01 and admixture proportions between 0 and 1) are shown for each target population.

| target | chisq | p | Yukon_Pleistocene | Central_AB_4k | wood_Late_Holocene | historical_WBNP | Bighorn_WY_250 |
|---|---|---|---|---|---|---|---|
| WBNP | 2.473 | 0.116 | - | - | - | 0.776 | 0.224 |
| ELKw | 3.922 | 0.048 | - | 0.122 | - | 0.878 | - |
| ELKw | 2.508 | 0.113 | - | - | - | 0.882 | 0.118 |
| MBS | 0.223 | 0.637 | - | - | 0.534 | 0.466 | - |
| MBS | 0.480 | 0.489 | - | - | - | 0.799 | 0.201 |
| RLBH | 1.973 | 0.160 | - | - | - | 0.594 | 0.406 |
| WAB | 0.609 | 0.435 | - | - | 0.626 | - | 0.374 |
| WAB | 2.381 | 0.123 | - | - | - | 0.287 | 0.713 |

**Table S8. Individuals used for bison-cattle admixture source panels.**

| sample ID | accession | species | breed or population |
|---|---|---|---|
| angus1 | SAMN05216009 | taurine cattle | angus |
| angus2 | SAMN05216010 | taurine cattle | angus |
| angus3 | SAMN05216011 | taurine cattle | angus |
| angus4 | SAMN05216012 | taurine cattle | angus |
| angus5 | SAMN05216013 | taurine cattle | angus |
| angus6 | SAMN05216014 | taurine cattle | angus |
| dominette | SAMN03145444 | taurine cattle | hereford |
| hereford1 | SAMN05216015 | taurine cattle | hereford |
| hereford2 | SAMN05216016 | taurine cattle | hereford |
| hereford3 | SAMN05216017 | taurine cattle | hereford |
| hereford4 | SAMN05216018 | taurine cattle | hereford |
| hereford5 | SAMN05216019 | taurine cattle | hereford |
| hereford6 | SAMN05216020 | taurine cattle | hereford |
| shorthorn1 | SAMN05216071 | taurine cattle | shorthorn |
| shorthorn2 | SAMN05216072 | taurine cattle | shorthorn |
| simmental1 | SAMN05216027 | taurine cattle | simmental |
| simmental2 | SAMN05216028 | taurine cattle | simmental |
| simmental3 | SAMN05216029 | taurine cattle | simmental |
| simmental4 | SAMN05216030 | taurine cattle | simmental |
| simmental5 | SAMN05216031 | taurine cattle | simmental |
| SC14.PH043 | F-3246 | steppe bison | Siberia_Holocene |
| UP10.MS071 | YG 303.666 | steppe bison | Northern_Pleistocene |
| SC14.AE009 | F-3006 | steppe bison | Siberia_Pleistocene |
| SC13.MS118 | YG 403.148 | steppe bison | Northern_Pleistocene |

14. R. O. Stephenson, S. C. Gerlach, R. D. Guthrie, C. R. Harington, R. O. Mills, G. Hare, "Wood bison in late Holocene Alaska and adjacent Canada: paleontological, archaeological and historical records" in *People and wildlife in northern North America: Essays in honor*

*of R. Dale Guthrie*, S. C. Gerlach, M. S. Murray, Eds. (BAR International Series 944, Archaeopress, 2001). pp. 124-158.

15. Environment and Climate Change Canada, "Recovery strategy for the wood bison (*Bison bison athabascae*) in Canada" (Species at Risk Act Recovery Strategy Series, Environment and Climate Change Canada, Ottawa, 2018). viii + 59 pp.

39. K. Wang, L. Wang, J. A. Lenstra, J. Jian, Y. Yang, Q. Hu, D. Lai, Q. Qiu, T. Ma, Z. Du, R.

Abbott, J. Liu, The genome sequence of the wisent (*Bison bonasus*). *Gigascience* **6**, 1–5 (2017).

40. A. Bergström, L. Frantz, R. Schmidt, E. Ersmark, O. Lebrasseur, L. Girdland-Flink, A. T. Lin, J. Storå, K.-G. Sjögren, D. Anthony, E. Antipina, S. Amiri, G. Bar-Oz, V. I. Bazaliiskii, J. Bulatović, D. Brown, A. Carmagnini, T. Davy, S. Fedorov, I. Fiore, D. Fulton, M. Germonpré, J. Haile, E. K. Irving-Pease, A. Jamieson, L. Janssens, I. Kirillova, L. K. Horwitz, J. Kuzmanovic-Cvetković, Y. Kuzmin, R. J. Losey, D. L. Dizdar, M. Mashkour, M. Novak, V. Onar, D. Orton, M. Pasarić, M. Radivojević, D. Rajković, B. Roberts, H. Ryan, M. Sablin, F. Shidlovskiy, I. Stojanović, A. Tagliacozzo, K. Trantalidou, I. Ullén, A. Villaluenga, P. Wapnish, K. Dobney, A. Götherström, A. Linderholm, L. Dalén, R. Pinhasi, G. Larson, P. Skoglund, Origins and genetic legacy of prehistoric dogs. *Science* **370**, 557–564 (2020).

41. R. E. Vance, D. Emerson, T. Habgood, A mid-Holocene record of vegetative change in central Alberta. *Can. J. Earth Sci.* **20**, 364–376 (1983).

42. I. D. Campbell, C. Campbell, Late Holocene vegetation and fire history at the southern boreal forest margin in Alberta, Canada. *Palaeogeogr. Palaeoclimatol. Palaeoecol.* **164**, 263–280 (2000).

43. COSEWIC, "COSEWIC assessment and status report on the Plains Bison *Bison bison bison* and the Wood Bison *Bison bison athabascae* in Canada" (Committee on the Status of Endangered Wildlife in Canada, Ottawa, 2013). xv + 109 pp.

54. Y. Running Horse Collin, C. P. Bataille, S. Hershauer, M. Hunska Tašunke Icu, A. Nujipi, W. Justin, J. Stelkia, J. A. Stelkia, S. A. Topkok, B. G. Leonard, B. Soop, M. Gonzalez, A. Luta Wiŋ, W. Wiŋ, T. Omniya, B. Dull Knife, B. Means, C. Tecumseh Collin, M. Koskey, J. D. Kapp, Z. Landry, D. Fraser, J. Southon, E. E. Lindroos, A. Hassler, L. Chauvey, G. Tressières, L. Tonasso-Calvière, S. Schiavinato, A. Seguin-Orlando, A. Perdereau, P. H. Oliveira, J.-M. Aury, P. Wincker, I. V. Kirillova, S. K. Vasiliev, M. A. Kusliy, A. S. Graphodatsky, A. A. Tishkin, I. Barnes, P. Druckenmiller, C. N. Jass, R. D. E. MacPhee, C. I. Barrón-Ortiz, P. Groves, D. Mann, D. G. Froese, M. Wooller, J. H. Miller, B. Crowley, G. Zazula, E. Hall, S. Hewitson, B. Shapiro, L. Orlando, Sustainability insights from Late Pleistocene climate change and horse migration patterns. *Science* **388**, 748–755 (2025).

55. G. C. Frison, R. C. Mainfort, Eds., *Archeological and Bioarcheological Resources of the Northern Plains* (Arkansas Archeological Survey, 1996).

56. P. J. White, R. L. Wallen, Yellowstone bison—should we preserve artificial population substructure or rely on ecological processes? *J. Hered.* **103**, 751–3 (2012).

57. E. Santiago, C. Köpke, A. Caballero, Accounting for population structure and data quality in demographic inference with linkage disequilibrium methods. *Nat. Commun.* **16**, 6054 (2025).

58. J. M. Martin, J. I. Mead, P. S. Barboza, Bison body size and climate change. *Ecol. Evol.* **8**, 4564–4574 (2018).

59. V. Geist, *Buffalo Nation: History and Legend of the North American Bison* (Voyageur Press, Stillwater, OK, 1998).

68. R. D. Guthrie, *Frozen Fauna of the Mammoth Steppe* (University of Chicago Press, Chicago, IL, 1989).

74. C. E. Schweger, M. Hickman, Holocene paleohydrology of central Alberta: testing the

general-circulation-model climate simulations. *Can. J. Earth Sci.* **26**, 1826–1833 (1989).

84. W. Olson, J. Janelle, *The Ecological Buffalo* (University of Regina Press, Regina, SK, Canada, 2022).

103. P. J. Reimer, W. E. N. Austin, E. Bard, A. Bayliss, P. G. Blackwell, C. Bronk Ramsey, M. Butzin, H. Cheng, R. L. Edwards, M. Friedrich, P. M. Grootes, T. P. Guilderson, I. Hajdas, T. J. Heaton, A. G. Hogg, K. A. Hughen, B. Kromer, S. W. Manning, R. Muscheler, J. G. Palmer, C. Pearson, J. van der Plicht, R. W. Reimer, D. A. Richards, E. M. Scott, J. R. Southon, C. S. M. Turney, L. Wacker, F. Adolphi, U. Büntgen, M. Capano, S. M. Fahrni, A. Fogtmann-Schulz, R. Friedrich, P. Köhler, S. Kudsk, F. Miyake, J. Olsen, F. Reinig, M. Sakamoto, A. Sookdeo, S. Talamo, The IntCal20 Northern hemisphere radiocarbon age calibration curve (0–55 cal kBP). *Radiocarbon* **62**, 725–757 (2020).

104. C. Bronk Ramsey, Bayesian analysis of radiocarbon dates. *Radiocarbon* **51**, 337–360 (2009).

105. K. Wang, J. A. Lenstra, L. Liu, Q. Hu, T. Ma, Q. Qiu, J. Liu, Incomplete lineage sorting rather than hybridization explains the inconsistent phylogeny of the wisent. *Commun. Biol* **1**, 169 (2018).

106. M. Gautier, K. Moazami-Goudarzi, H. Levéziel, H. Parinello, C. Grohs, S. Rialle, R. Kowalczyk, L. Flori, Deciphering the wisent demographic and adaptive histories from individual whole-genome sequences. *Mol. Biol. Evol.* **33**, 2801–2814 (2016).

107. K. Wecek, S. Hartmann, J. L. A. Paijmans, U. Taron, G. Xenikoudakis, J. A. Cahill, P. D. Heintzman, B. Shapiro, G. Baryshnikov, A. N. Bunevich, J. J. Crees, R. Dobosz, N. Manaserian, H. Okarma, M. Tokarska, S. T. Turvey, J. M. Wójcik, W. Zyla, J. M. Szymura, M. Hofreiter, A. Barlow, Complex admixture preceded and followed the extinction of wisent in the wild. *Mol. Biol. Evol.* **34**, 598–612 (2017).

108. M. P. Verdugo, V. E. Mullin, A. Scheu, V. Mattiangeli, K. G. Daly, P. Maisano Delser, A. J. Hare, J. Burger, M. J. Collins, R. Kehati, P. Hesse, D. Fulton, E. W. Sauer, F. A. Mohaseb, H. Davoudi, R. Khazaeli, J. Lhuillier, C. Rapin, S. Ebrahimi, M. Khasanov, S. M. F. Vahidi, D. E. MacHugh, O. Ertuğrul, C. Koukouli-Chrysanthaki, A. Sampson, G. Kazantzis, I. Kontopoulos, J. Bulatovic, I. Stojanović, A. Mikdad, N. Benecke, J. Linstädter, M. Sablin, R. Bendrey, L. Gourichon, B. S. Arbuckle, M. Mashkour, D. Orton, L. K. Horwitz, M. D. Teasdale, D. G. Bradley, Ancient cattle genomics, origins, and rapid turnover in the Fertile Crescent. *Science* **365**, 173–176 (2019).

136. G. C. Frison, L. C. Todd, Eds., *Horner Site* (Elsevier Science, 2014).

137. E. B. Howard, The Finley site: Discovery of Yuma points, in situ, near Eden, Wyoming. *Am. Antiq.* **8**, 224–234 (1943).

138. G. C. Frison, The Carter/Kerr-McGee Paleoindian site: Cultural resource management and archaeological research. *Am. Antiq.* **49**, 288–314 (1984).

139. R. Maier, P. Flegontov, O. Flegontova, U. Işıldak, P. Changmai, D. Reich, On the limits of fitting complex models of population history to f-statistics. *eLife* **12**, e85492 (2023).

140. G. Renaud, K. Hanghøj, T. S. Korneliussen, E. Willerslev, L. Orlando, Joint estimates of heterozygosity and runs of homozygosity for modern and ancient samples. *Genetics* **212**, 587–614 (2019).

141. W. Haak, I. Lazaridis, N. Patterson, N. Rohland, S. Mallick, B. Llamas, G. Brandt, S. Nordenfelt, E. Harney, K. Stewardson, Q. Fu, A. Mittnik, E. Bánffy, C. Economou, M. Francken, S. Friederich, R. G. Pena, F. Hallgren, V. Khartanovich, A. Khokhlov, M. Kunst, P. Kuznetsov, H. Meller, O. Mochalov, V. Moiseyev, N. Nicklisch, S. L. Pichler, R. Risch, M. A. Rojo Guerra, C. Roth, A. Szécsényi-Nagy, J. Wahl, M. Meyer, J. Krause, D. Brown, D. Anthony, A. Cooper, K. W. Alt, D. Reich, Massive migration from the steppe was a source for Indo-European languages in Europe. *Nature* **522**, 207–211 (2015).

142. P. Skoglund, J. C. Thompson, M. E. Prendergast, A. Mittnik, K. Sirak, M. Hajdinjak, T. Salie, N. Rohland, S. Mallick, A. Peltzer, A. Heinze, I. Olalde, M. Ferry, E. Harney, M. Michel, K. Stewardson, J. I. Cerezo-Román, C. Chiumia, A. Crowther, E. Gomani-Chindebvu, A. O. Gidna, K. M. Grillo, I. T. Helenius, G. Hellenthal, R. Helm, M. Horton, S. López, A. Z. P. Mabulla, J. Parkington, C. Shipton, M. G. Thomas, R. Tibesasa, M. Welling, V. M. Hayes, D. J. Kennett, R. Ramesar, M. Meyer, S. Pääbo, N. Patterson, A. G. Morris, N. Boivin, R. Pinhasi, J. Krause, D. Reich, Reconstructing prehistoric African population structure. *Cell* **171**, 59–71.e21 (2017).
